# Desiccation-driven senescence and its repression in *Xerophyta schlechteri* are regulated at extremely low water contents

**DOI:** 10.1101/2021.06.04.447054

**Authors:** Astrid L. Radermacher, Brett Williams, Arash Iranzadeh, Halford Dace, Sagadevan Mundree, Henk W.M. Hilhorst, Jill M. Farrant

## Abstract

- Vegetative desiccation tolerance, the ability to survive loss of over 90% of cellular water, is an extremely rare trait in Angiosperms. *Xerophyta schlechteri* survives such extreme water deficit by entering prolonged quiescence and suppressing drought-induced senescence in most of the leaf area, except the apical tip. Information on the molecular regulation of senescence in such plants is scarce and this is the first study to investigate such regulation in senescing and non-senescing tissues of the same leaf.
- Genome-wide RNA sequencing enabled comparison of senescent and non-senescent tissues during desiccation and early rehydration, establishment of the water content range in which senescence is initiated and identification of molecular mechanisms employed to bring about cellular death.
- Senescence-associated genes (*Xs*SAG) specific to this species were identified and two potential regulatory sites were enriched in regions upstream to these *Xs*SAGs, allowing us to create a model of senescence regulation in *X. schlechteri* based on homology with known Arabidopsis senescence regulators.
- We hypothesise that desiccation-driven senescence occurs as a result of a convergence of signals around MAPK6 to trigger WRKY-mediated ethylene synthesis and *Xs*SAG expression, not unlike aging and stress-related senescence in Arabidopsis, but at remarkably lower water contents (<35% RWC).

## Introduction

Vegetative desiccation tolerance, the ability to survive loss of over 90% of cellular water, is rare in Angiosperms (0.04% of species), these collectively being termed resurrection plants. While considerable molecular data is available on mechanisms that enable survival of extreme water loss (for recent reviews, see Gariola *et al*., 2017; Oliver *et al*., 2020), there is little information yet on regulation of senescence in such plants. It has been postulated that resurrection plants are able to prevent cellular death processes during desiccation, through deployment of molecular protectants (anti-oxidants, LEAs, chaperones), clearing of debris (unfolded protein response and autophagy) as well as attenuation of cellular death regulators (e.g. Senescence-associated receptor kinase (SIRK) and the transcription factor WRKY53) (Griffiths *et al*., 2014; Willams *et al*., 2015). However, to date no comparative studies have been performed on surviving compared with senescing tissues of individual species of resurrection plants, and thus no direct mechanisms for transcriptional regulation thereof have been identified.

*Xerophyta schlechteri* (syn. *Xerophyta viscosa*) is a valuable species in which to study the phenomenon of senescence prevention in resurrection plants. Its survival of desiccation has been well characterised at the physiological and molecular levels (Sherwin and Farrant, 1996; 1998; Mundree and Farrant, 2000; Farrant *et al*., 2015) and its genome has been sequenced (Costa *et al*., 2017a). This poikilochlorophyllous monocot breaks down most of its chlorophyll during drying but represses senescence in the bulk of its tissues, with only the leaf apices, the oldest tissue, succumbing to senescence during desiccation (Radermacher *et al*., 2019). This process is likely initiated below 35% relative water content (RWC), but the question as to when during drying senescence-associated gene (*Xs*SAGs) expression is transcriptionally activated remains unanswered. A comparative transcriptomic approach investigating gene expression in tissue destined for senescence (senescent tissue, ST) with that of non-senescent tissue (NST), will enable assessment of mechanisms supressing senescence in the latter.

Recent studies have demonstrated the power of network analysis in addition to conventional differential gene expression analysis in that it provides insight into how networks of genes and gene products interact to enable plant survival under stressful conditions (Costa *et al*., 2017a; Liu *et al*., 2019; Woo *et al*., 2019). This is particularly true in *X. schlechteri* where the genome is octoploid and many (often duplicated) genes work in concert to bring about various cellular and whole-plant changes, resulting in ultimate desiccation tolerance and quiescence (Costa *et al*., 2017a). Such analysis involves, *inter alia*, deployment of techniques such as K-Means clustering and co-expression network analysis, where the expression patterns of genes are grouped based on the similarity of their temporal expression. We utilized these tools to describe the senescence-associated transcriptome (Senescome) of *Xerophyta schlechteri*, in the context of desiccation tolerance (Desiccome), to determine when this subset of *Xs*SAGs is expressed in a drying cycle, identify potential regulators thereof and describe the cellular context for senescence initiation. We postulate how this process may be supressed in NST.

## Materials and Methods

### Plant material and sampling procedures

Plants were accessed, maintained and subject to dehydration and recovery treatments as described in Radermacher *et al*., (2019). Leaf tissue (10 leaves per plant) was sampled daily one hour after dawn. Each leaf tissue was dissected into an apical region (3cm) designated as ST, a region of 1cm below this, termed pre-senescent tissue (PST: that which will survive a first dehydration but will senesce in a future desiccation event) and a section of 1cm, cut from the remaining NST (Fig. **1a**). These were cut in half down the midrib, with one half being frozen in liquid nitrogen and stored for RNA extraction and the other half used for absolute and RWC determination. RWC was determined as described in Radermacher *et al*., (2019). As plants, and leaves within a single plant, dehydrate at different rates, ultimately three biological replicates, each corresponding to a designated water content (outlined below) were used for RNA extraction. Tissue utilized from fully hydrated plants, prior to dehydration, included only the NST and ST samples. Thus, 51 samples were prepared for RNA extraction.

**Figure 1:**
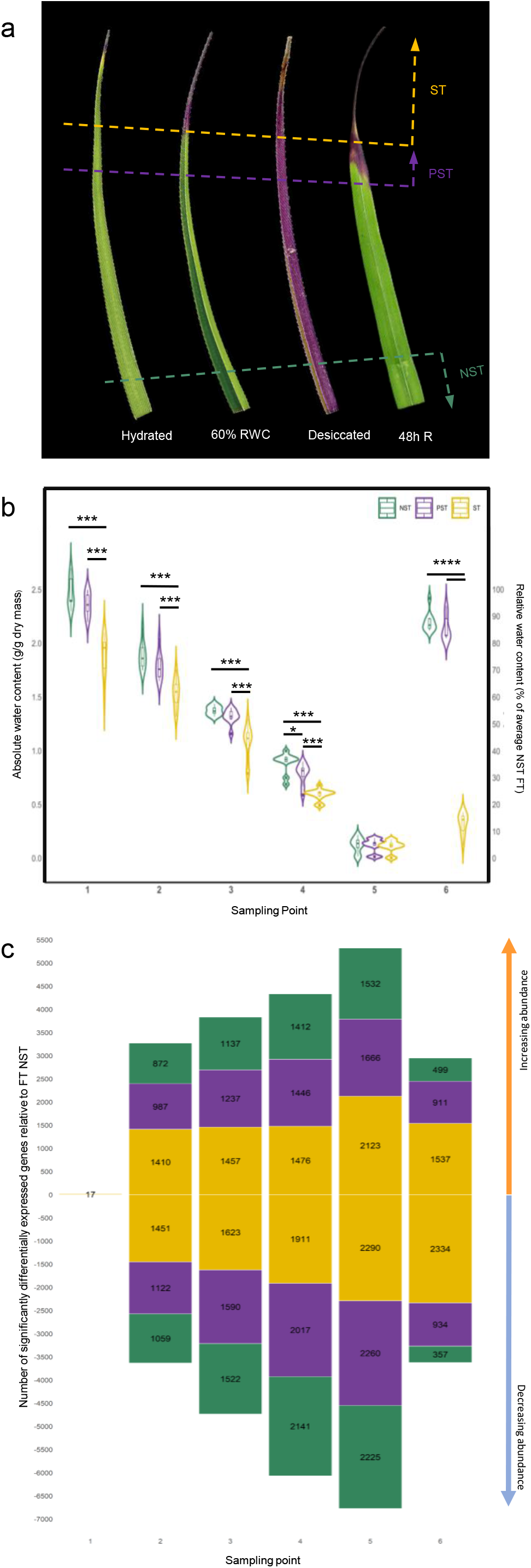
Sampling protocol and changes in tissue appearance, water content and differential gene expression during desiccation and rehydration. **a**. Typical appearance of leaves at full turgor, partially dehydrated, desiccated and partially rehydrated (48h) states. Samples with a similar RWC value in NST (green zone) were chosen and grouped into sampling points for RNA extraction, utilizing the ST (yellow zone) and PST (purple zone) of the same leaves despite variability in RWC. Sampling of leaves for transcriptomics was performed based on NST RWC values. **b**. Sampling point 1 represents the well-watered “full turgor” condition. Samples 2-4 represent drought stressed plants in descending order of severity (75%, 55% and 35% of the average RWC in NST). Sampling point 5, where the absolute water content between tissues is not significantly different, represents the desiccated state. Sampling point 6 represents 48h post rehydration. The variability of RWC is shown here, with the results of a Wilcoxon test for each tissue type in the same sampling point given (only significantly differences are shown). Significance codes: “****” corrected pval < 0,0001; “***” corrected pval < 0,001; “**” corrected pval < 0,01; “*” corrected pval < 0,05. **c**. DEGs where the number of genes with a significant increase in expression is shown on the positive y axis and the number of genes with a significant decrease in expression is shown on the negative y axis. Genes are considered significantly differentially expressed if they exhibit a log2 fold change value greater than 2 or less than -2, compared to control and an Exact test corrected pval <0.05 (C).

### RNA extraction from *X. schlechteri*

Total RNA was extracted from hydrated, dehydrating (75%, 55%, 35%), desiccated (5% relative water content (RWC)) and rehydrated (48h – varying RWC values) leaf tissues of *X*.*schlechteri*. A modified extraction method was used, using Tri-reagent (Sigma-Aldrich Corporation, St. Louis, MO, USA) and the RNEasy mini-kit (Qiagen, Hilden, Germany). Samples held in storage at-80°C were transferred under liquid nitrogen into sterilized 2mL Eppendorff tubes containing three stainless steel bearings (pre-treated by soaking in chloroform and autoclaved twice along with the tubes). Before the tissue was allowed to thaw, 1mL cold Tri-reagent was pipetted into the tubes, ensuring that all tissue was completely submerged. The samples were maintained on ice until being transferred to an in-house homogenizer, where they were homogenized at room temperature for 12 minutes. The samples were placed on ice and 200µl chloroform was added to each tube. The samples were mixed by inversion and maintained on ice for 2 min. The tubes were then centrifuged at 13000g for 15 min at 4°C. The upper aqueous phase of each sample was transferred to the RNEasy mini-kit Shredder column and the manufacturer’s instructions for RNA extraction were followed with the following modifications: the RW1 buffer wash step was repeated with a volume of 200µl, the RPE buffer wash step was repeated twice and the RNA was eluted in RNase-free water pre-heated to 55°C after incubating for 10 min at room temperature. RNA integrity and quality were verified using a Bioanalyzer (Agilent technologies, USA).

### High-throughput sequencing and transcriptome annotation

Library preparation was conducted using the Illumina RNA-seq kit (San Diego, California) according to the manufacturer’s instructions. The resulting cDNA libraries were sequenced using the Illumina MiSeq500 (San Diego, California) platform at Queensland University of Technology’s Central Analytical Research Facility. The Tuxedo suite (Trapnell *et al*., 2014) was used for annotation and mapping to the reference genome for this species (Costa *et al*., 2017a). The genome indexing was built by bowtie2. The reference transcriptome indexing and the alignment of the RNA-seq reads to the reference genome were performed by tophat. The transcriptome assembly of samples was performed by cufflink and the assembled transcriptome of different samples were merged by cuffmerge (Trapnell *et al*., 2014). Differential gene expression was performed using cuffdiff on FPKM (fragments per kilobase of exon per million mapped reads) gene expression data. Genes were considered differentially expressed if they exhibited a log2 fold change value greater than 2, compared to control and Exact test corrected pval <0.05. Two datasets are presented: the Desiccome (those genes which are significantly differentially expressed in all tissues during water deficit stress) and the Senescome (those genes differentially expressed in the airdry condition in ST, relative to AD NST). These data were plotted as a series of heatmaps in R (version 3.5.1; R Development Core Team, 2011) based on their Mercator Bins (Lohse *et al*., 2014) using the packages ggplot2 (Wickham, 2016) and wesanderson, and tabulated using the package dataTables (Jardine, 2019) and sparkline (Vaidyanathan, 2016). These plots and tables were rendered as interactive apps with the R package Shiny by RStudio (RStudio Team, 2015). The same method of analysis and data visualisation was followed for the Senescome, where genes significantly accumulated in AD ST relative to AD NST are visualised. The apps are available at astridite.shinyapps.io/XeroDesiccome and at astridite.shinyapps.io/XeroSenescome. The SwissProt, Gene Ontology and KEGG annotations were provided with the genome (Costa *et al*., 2017a). Additional annotation was performed by Mercator (Lohse *et al*., 2014) to provide plant-specific “BIN” ontologies.

### qPCR validation of transcriptome

Ten DEGs were selected from the transcriptomic analysis for validation using qPCR. The RNA extracted (1 µg per sample) for the transcriptomic analysis from all three tissues; 100%, 55% RWC and 48h RH, was converted to cDNA using Phusion HiFi Taq polymerase as per the manufacturer’s instructions. qPCR was performed on 2µl of the resulting cDNA. The primers and the amplification efficiency of the ten DEGs were designed and optimised by assessing the relative transcript abundance of each GOI through RT-qPCR. Pipetting of the reaction mix (5µl SYBR green, 0.3µl forward and reverse primer, 2µl cDNA, 2.4µl nuclease-free water) was performed using an Eppendorf epMotion® 5075 automated pipetting machine. The amplification profile was as follows: 30 cycles of 15 sec at 95°C, followed by 60 sec at 60°C. Expression was normalised using PolyUbiquitin10 as the reference gene and log2FC values were calculated based on comparison to the FT NST condition. The resulting comparative plot (Fig. **S6**) was created in RStudio using the ggplot2 package.

### Gene expression profiling

To explore and describe the temporal trends in the dataset, twelve K-means clusters were calculated in Genesis 1.8.1 from gene expression (Fragments per kilobase per million mapped reads - FPKM) data divided into the NST, PST and ST tissue types, which were clustered based on gene-level normalisation (Sturn et al., 2002). Further to this end, Pearson correlation coefficients were calculated between each gene pair to yield a correlation matrix in which each gene was compared to all genes in the dataset to determine co-expression (defined as R^2^ >0.94). Co-expressed genes were visualised in Cytoscape v3.7.1, where distances between nodes (edges) were representative of the level of co-expression over a threshold of 0.94; the higher the correlation, the closer the nodes. The network was displayed using the yGraph Organic layout and coloured based on the K-means cluster, degree and the differential gene expression results.

### Promoter motif enrichment analysis

Enriched promoter regions in the upstream regions (up to 1kb of sequence) of the senescence-specific subcluster of the gene expression network were identified using DREME from the MEME suite of promoter analysis tools (Bailey *et al*., 2009; Bailey, 2011) with the upstream regions of the entire network as a control. Significantly enriched sequences were calculated using Fischer’s exact test (p < 0.05). Transcription factors that bind to these enriched promoters were identified using TomTom (Jaspar Non-redundant 2018 plants database) (Gupta *et al*., 2007) and annotated with *X. schlechteri* genomic identifiers in Microsoft Excel 365. The frequency of binding site occurrences was studied using Centrimo from the MEME suite (Bailey & Machanick, 2012). Signalling pathway construction was performed using these annotations by incorporating transcription factor and signalling kinase nodes from the network as well as signalling information obtained from the cited literature.

### ABA quantification by UHPLC–MS/MS

Freeze-dried samples of 1mg were ground and extracted as described (Floková *et al*. 2014), with the following adjustments: only deuterated ABA was used as an internal standard and prior to injection, samples were filtered to remove particulates

## Results

### Desiccation-driven senescence

With each desiccation event, minimal tissue from the apex of *X. schlechteri* leaves does not recover. This senescent region (ST) is bordered by a purple zone destined to senesce at the next desiccation event (pre-senescent tissue, PST) (Fig. **1a**). This failure of apical tissue to recover after rehydration is observed regardless of the degree of leaf expansion, with all leaves exhibiting the same pattern during and after a desiccation event. Only fully expanded leaves that had never undergone a desiccation event were studied.

### Relative water content and differential gene expression

Samples for RNA extraction were grouped based on their RWC following water withholding (Fig. **1b**). As previously reported for this species (Radermacher *et al*., 2019), the RWC of sampled ST was always lower than for NST, except in the desiccated state, when all tissues come into equilibrium. Examination of the PST was undertaken to determine whether there existed a spatial progression towards the senescent state. There was no significant difference in RWC between NST and PST tissues in rate and totality of dehydration, nor in the ability to rehydrate fully. Forty-eight hours after rehydration, NST and PST reached an average RWC of approximately 90%, whereas ST reached only 18% (Fig. **1b**). In the hydrated state only 17 genes were differentially expressed between NST and ST (Fig. **1c**). As the NST dehydrated to 75% RWC, over 1800 differentially expressed genes (DEGs) were observed for each tissue type (relative to hydrated NST). When the NST was at 35% RWC, there was very little difference in the number of DEGs that were expressed between tissue types (Fig. **1c**). However, when the plants desiccated towards the airdry state (<10% RWC), the difference in transcript abundance between tissues became evident, with 591 more DEGs upregulated in ST than in its NST counterpart. Upon rehydration, the number of DEGs in NST and PST declined but the number of DEGs in ST remained elevated (Fig. **1c**).

### K-means Clustering and Co-expression Network

Twelve K-means clusters were organized into subcategories (“superclusters”) based on their responses during desiccation (Fig. **2**). Clusters 8, 10 and 12 were designated the reduced and recovered supercluster (RRS) based on their relative diminishment in all three tissues during drying with recovery evident on rehydration in only NST and PST. Clusters 1 and 9 were grouped into the early response supercluster (ERS) based on their early (75%-55% RWC) accumulation and clusters 4, 5 and 7 were grouped into the late response supercluster (LRS) based on their late accumulation (<35% RWC). Cluster 2 (drought and desiccation cluster - DDC) represents those transcripts accumulated at the onset of water deficit stress, which remain elevated throughout drying and desiccation. These expression patterns are consistent with the early and late transcriptome responses described by Costa *et al*. (2017a). Clusters 3, 6 and 11 represent transcripts which were elevated in ST in late stages of drying relative to NST and PST, thus grouped into a late response ST (LR-ST) supercluster (See supplementary data for enriched GO categories of each cluster). A Pearson correlation network was created to further explore relationships between genes that cluster based on their expression pattern, and to cross-validate the K-means clustering result with another unsupervised learning technique (Fig. **2b**). The spatial separation of these clusters closely aligned with their associated K-means supercluster (Fig. **2a**), allowing later dissection of the senescence-specific subcluster away from the global network, and for downstream regulation modelling (Figs **6** and **7**).

**Figure 2:**
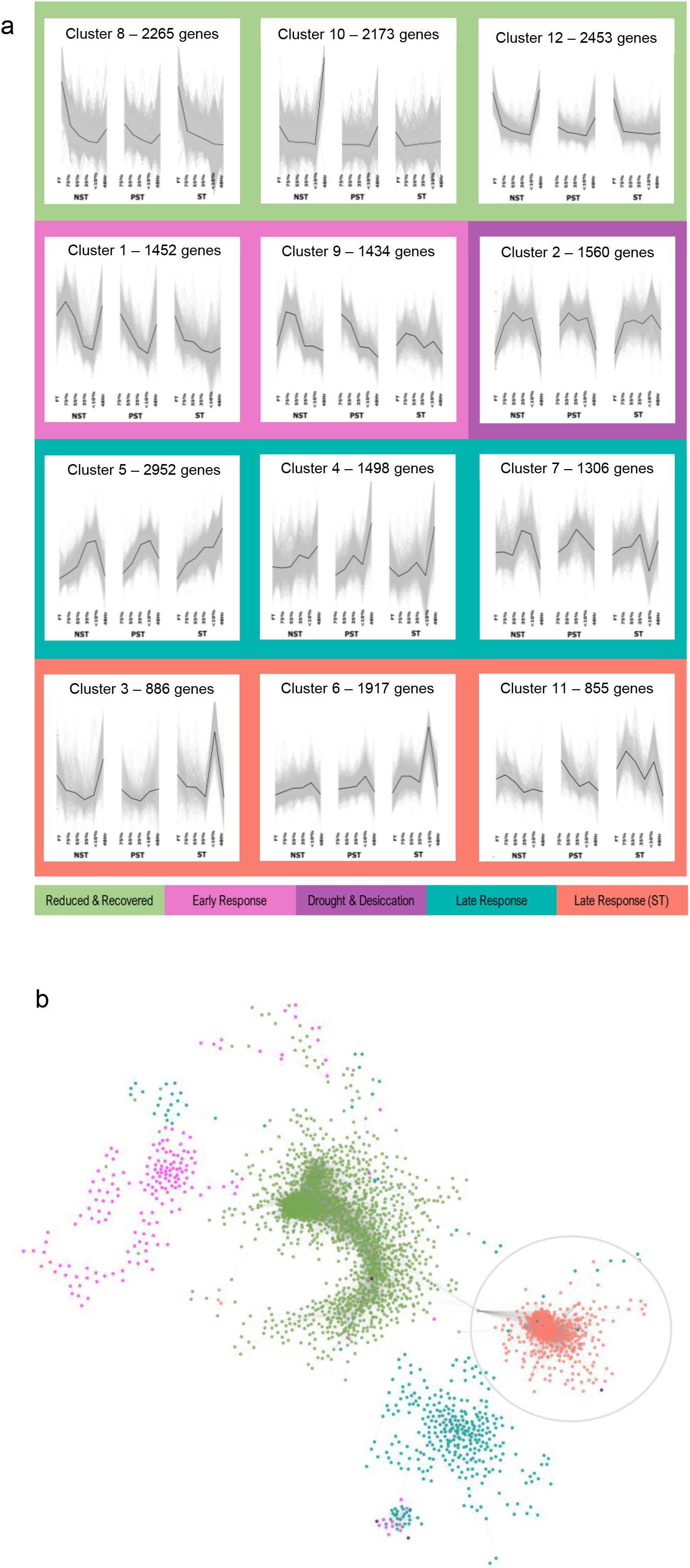
K-means Clustering and Co-expression Network Analysis. **a**. Transcript abundance (FPKM) assembled into 12 K-means clusters. The clusters are grouped into “superclusters” based on the response of to water deficit. Green represents those genes whose expression declines with water deficit and recovers upon rehydration. Pink represents those genes which peak in expression in NST and PST at 75% RWC. Purple represents those genes which are highly expressed in all three tissues from the onset of water stress until rehydration, when their expression declines. Genes in teal represent those who accumulate in the late stages of drying (<35% RWC) in NST and PST. Genes in orange represent those highly expressed in desiccated ST . **b**. A co-expression network of pearson-correlated (R2 >0.94) genes was generated and overlaid with these K-means clusters. The senescence-specific subnetwork (designated as such due to the high level of gene expression in desiccated ST) is circled in grey.

Based on clustering results, transcripts most likely to be associated with the onset of senescence were highly accumulated in the desiccated state. To explore the desiccation response independently of the senescence response, two subsets of the data were created; the Desiccome, which encompasses transcripts in the DDC, ERS and LRS clusters significantly differentially accumulated compared to FT NST; and the Senescome, which encompass transcripts highly expressed in desiccated ST relative to desiccated NST.

### Desiccome: expression of genes related to desiccation tolerance

This data set (explorable on the interactive Desiccome app: https://astridite.shinyapps.io/XeroDesiccome/) encompasses transcripts associated with vegetative desiccation tolerance in this species (Costa *et al*. (2017a). These included significant accumulation of, *inter alia*, late embryogenesis abundant (LEA) proteins, heat shock proteins, molecular chaperones and anti-oxidants during water deficit (Desiccome, Fig. **3**). In line with metabolomic studies in other resurrection species (Moyankova *et al*., 2014; Suguiyama *et al*., 2014; Yobi *et al*., 2017), transcripts associated with accumulation of raffinose family oligosaccharides (RFOs) and trehalose were significantly accumulated (Sugars tab, Desiccome). The abundance of lipid-degrading and -modifying enzyme transcripts increased during drying, as did those encoding oleosins (Lipids tab, Desiccome). Patterns of expression of signalling molecules (ABA, calcium, light, G-proteins and receptor kinases) was consistent between tissues (Signalling tab, Desiccome). ABA regulates abiotic stress and desiccation tolerance in this species (Costa *et al*., 2017a), and so the presence of ABA-responsive genes (ABI5, GEM-like protein 3 and several PP2Cs) is not unexpected. Serine, aspartate, cysteine and AAA-type protease genes were significantly upregulated during desiccation in all tissues, as were 38 E3 family ubiquitin ligase encoding genes. The upregulation of these genes may indicate functioning of the unfolded protein response (UPR), enabling degradation of misfolded proteins produced by the endoplasmic reticulum during dehydration (Ye *et al*., 2011; Griffiths, *et al*., 2014). This in turn could indicate activation of the ubiquitin proteasomal system as an early and less severe response compared to autophagy. Notably, two metacaspase 1 genes, encoding a caspase-like protein that is implicated in bringing about “superoxide-dependent cell death in a reactive oxygen-sensitized state” (Coll *et al*., 2014) were upregulated in the late stages of desiccation in all three tissue types (Protein tab, Desiccome). DNA repair transcripts (DNA repair protein UVH3, Deoxyribodipyrimidine photo-lyase and DNA-damage-repair/toleration protein DRT111 (DNA tab, Desiccome) were also upregulated in the late stages of desiccation in the NST and PST but not to the same degree in ST. Several transcriptional regulators significantly accumulated during dehydration include several MYB, DIVARICATA, heat stress, DREB and ZAT transcription factors. Interestingly, three transcripts homologous with ORE1, a known pro-senescence regulator in Arabidopsis (Matallana-Ramirez *et al*., 2013), were highly accumulated in all three tissues during desiccation (RNA tab, Desiccome).

**Figure 3:**
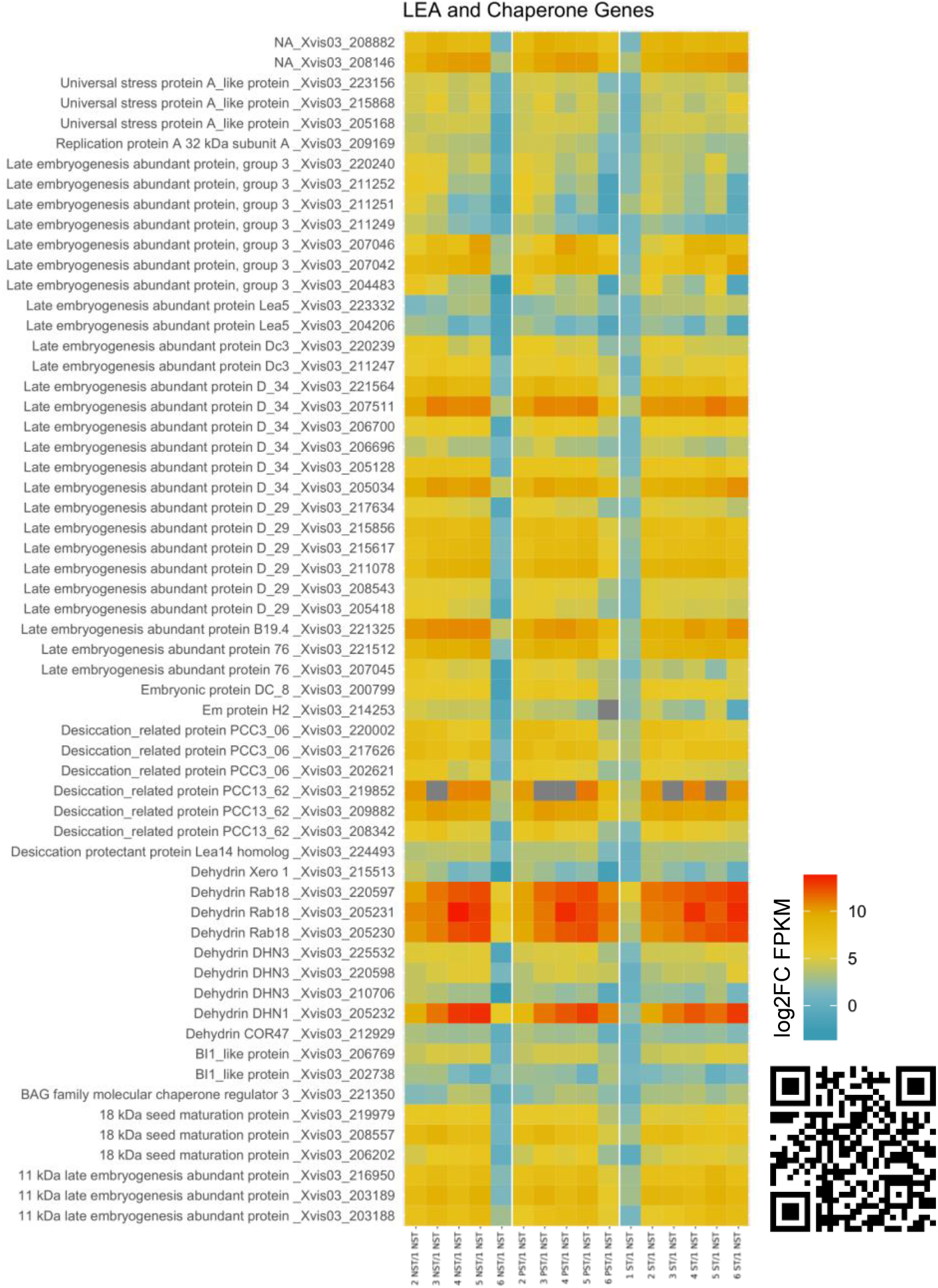
The Desiccome - desiccation-required genes are highly accumulated in all leaf tissues of *X. schlechteri*. Log2 fold change values relative to FT NST are shown in NST, PST and ST during water deficit and rehydration. Genes are annotated with their SwissProt annotation and genome ID. RWC values are representative of NST for easy comparison of tissues. White vertical lines separate tissue types. LEA and chaperone genes are shown here, but **t**he full Desiccome dataset is available at the QR code or at https://astridite.shinyapps.io/XeroDesiccome/

Senescent tissue did not deviate dramatically from NST and PST in terms of expression of these genes crucial for desiccation tolerance, suggesting that cellular death occurs as a programmed response to prolonged exposure to extreme stress in this tissue, rather than its inability to mitigate stress.

### Senescome: expression of desiccation-driven senescence genes

The interactive Senescome (https://astridite.shinyapps.io/XeroSenescome/) reports DEGs in desiccated ST relative to desiccated NST (senescence-related subset, Fig. **4**). Comparison of tissues within this sampling point enables dissection of senescence-related genes independently from desiccation-related genes (which are highly expressed in both tissues), as opposed to a comparison to fully hydrated NST, which would conflate senescence and desiccation tolerance processes.

**Figure 4:**
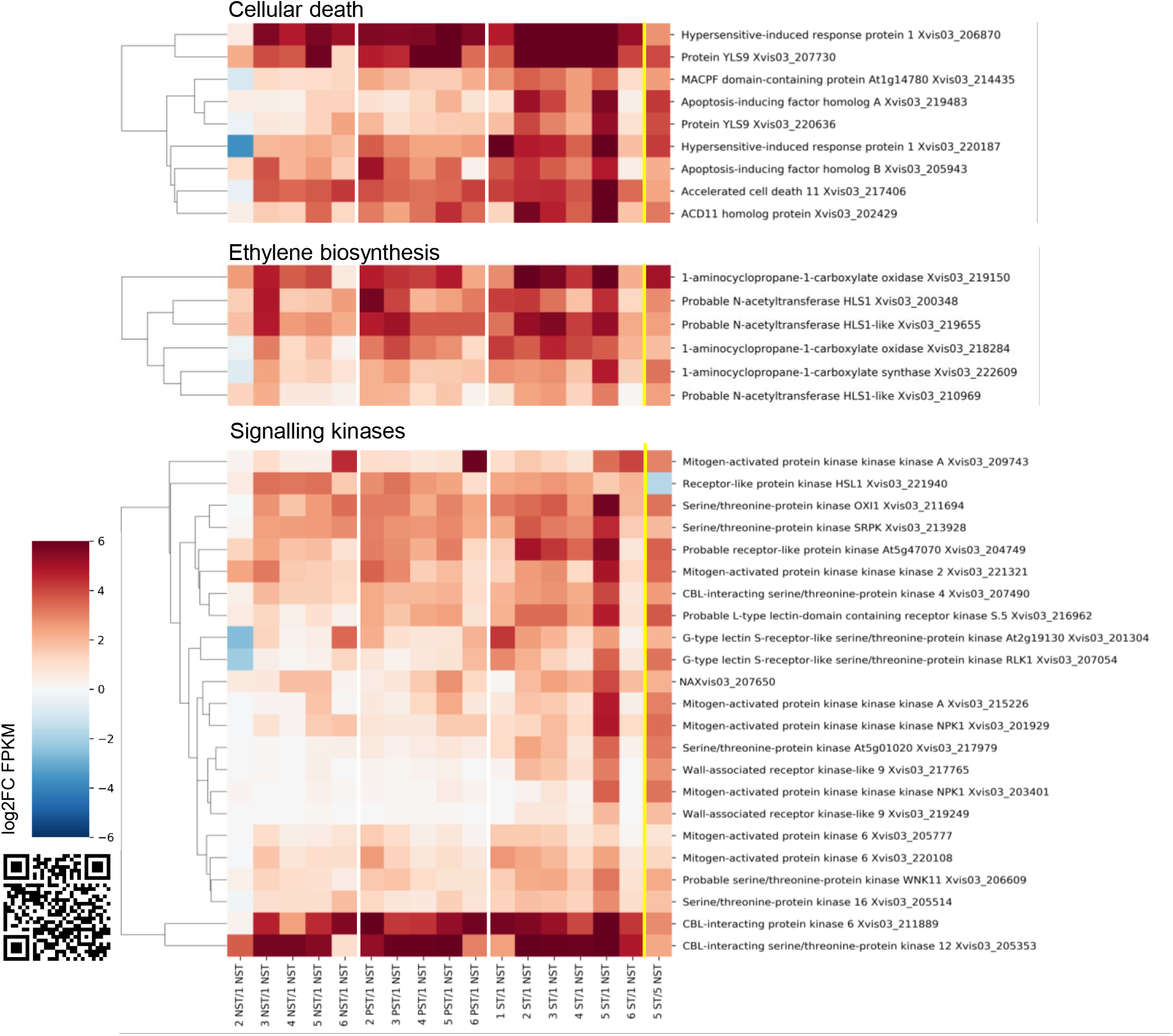
The Senescome – transcription of desiccation-driven senescence genes is most abundant below 35% RWC (sampling point 5) in apical ST. Senescence-specific DEGs were identified by comparing desiccated ST to desiccated NST. A subset of genes in the Mercator categories to do with senescence and signalling are shown here, with their Log2 fold change values relative to hydrated NST shown in NST, PST and ST during water deficit and rehydration. Genes are annotated with their SwissProt annotation and genome ID. X-axis labels are representative of the comparison, with the sampling point (number 1-6) and tissue type (NST/PST/ST) shown relative to hydrated NST (1 NST). White vertical lines separate tissue types (NST-PST-ST). The right-most column represents gene expression in desiccated ST relative to desiccated NST (5 ST/5 NST). The full Senescome dataset is available as a supplementary data set, or at the QR code (https://astridite.shinyapps.io/XeroSenescome/)

#### Cell death

Mercator-annotated cell death transcripts accumulated late in the desiccation timeline, and to a higher degree in ST. Transcripts encoding apoptosis-inducing factor homologues A and B, Yellow Leaf Specific proteins, accelerated cell death 11 and hypersensitive-induced response protein 1 were all highly expressed in response to desiccation in ST (Cell tab, Senescome app; Fig. **4**).

#### Hormone metabolism

Several transcripts coding for hormone synthesis and binding were significantly accumulated in desiccated ST relative to NST, most notably ethylene biosynthesis enzymes 1-aminocyclopropane-1-carboxylate (ACC) oxidase and ACC synthase (Signalling-Hormones tab, Senescome). The transcript encoding a gibberellin deactivating enzyme, gibberellin 2-beta-dioxygenase, was highly accumulated in this subset, indicating that gibberellin-responsive gene expression is tempered. Jasmonate synthesis may also be triggered, as transcripts encoding 12-oxophytodienoate reductase 1 and allene oxide synthase 1 and 2, role-players in oxylipin metabolism, were significantly accumulated.

#### Macromolecular degradation

Protein degradation is favored over protein synthesis in desiccated ST, with 40 transcripts indicating various protein degrading mechanisms being significantly differentially accumulated (Protein tab, Senescome; Fig. **5**). Those involved with proteasomal degradation were most abundant, as well as a number of aspartate, serine and metallo-proteases (also enriched during senescence in barley and Arabidopsis) (Parrott *et al*., 2007; Golldack et al., 2002). A cysteine protease vacuolar processing enzyme (VPE) was highly expressed in this dataset and has been shown to act during senescence (Hatsugai *et al*., 2015) by mediating vacuolar degradation and initiation of proteolysis and PCD. Protein degradation processes also included four members of the autophagosome complex: autophagy-related protein 8C, 13, 18f and 18h transcripts, indicating that the cells might be triggering autophagy, unlike the surviving tissues.

**Figure 5:**
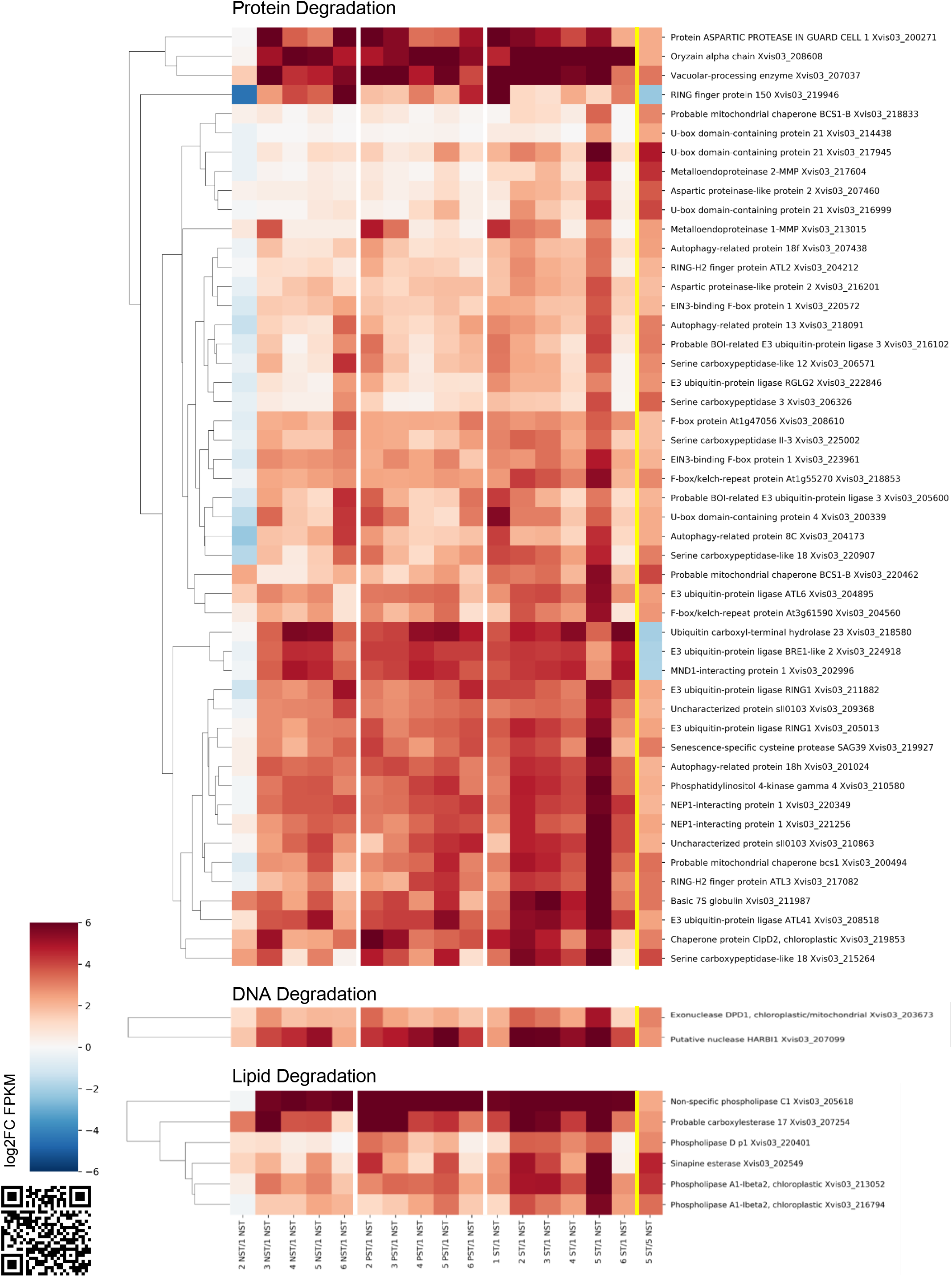
The Senescome – transcription of macromolecular degradation genes is most abundant below 35% RWC (sampling point 5) in apical ST. Senescence-specific DEGs were identified by comparing desiccated ST to desiccated NST. A subset of genes in the Mercator categories to do with senescence and signalling are shown here, with their Log2 fold change values relative to hydrated NST shown in NST, PST and ST during water deficit and rehydration. Genes are annotated with their SwissProt annotation and genome ID. X-axis labels are representative of the comparison, with the sampling point (number 1-6) and tissue type (NST/PST/ST) shown relative to hydrated NST (1 NST). White vertical lines separate tissue types (NST-PST-ST). The right-most column represents gene expression in desiccated ST relative to desiccated NST (5 ST/5 NST). The full Senescome dataset is available as a supplementary data set, or at the QR code (https://astridite.shinyapps.io/XeroSenescome/)

A number of DNA and RNA nucleases were upregulated in desiccated ST including an organelle-specific nuclease (DPD1) and ribonucleases (DNA tab, Senescome). Endonuclease 1 transcripts were highly also accumulated. This calcium-dependent senescence-specific nuclease degrades both DNA and RNA during the terminal stages of leaf senescence in Arabidopsis (Perez-Amador *et al*., 2000; Farage-Barhom *et al*., 2008). Additionally, structural maintenance of chromosomes proteins 1 and 3 (cell-cell division tab, Senescome) were diminished in desiccated ST, indicating that the structural integrity of chromatin is not prioritized in this tissue. Ribosome biogenesis transcripts are also significantly diminished in ST (Protein-synthesis tab, Senescome), indicating that storage of transcripts for translation upon rehydration is deprioritised. Lipid degrading enzymes were highly upregulated in the Senescome, including phospholipases A1, C1 and D (Lipids tab, Senescome; Fig. **5**), all of which have been shown to accumulate in senescent Arabidopsis leaves (Schmid *et al*., 2005; Van Der Graaff *et al*., 2006; Breeze *et al*., 2011). Synthesis of phosphatidic acid, an important abiotic stress signalling lipid, is driven by the sequential action of phospholipases C or Dp1 and diacylglycerol kinase, all upregulated in ST. The appearance of four transcripts binned under sphingolipid biosynthesis is of interest, as this lipid family has been proposed as a regulator of apoptosis-like PCD in Arabidopsis (Alden *et al*., 2011) (Lipids tab, Senescome). Degradation of lipids may also be used to drive central metabolism for generation of NADH and ATP (Troncoso-Ponce *et al*., 2013) (Lipids tab, Senescome).

#### Metabolism: energy generation via alternate pathways and electron transport

Transcripts encoding glycolysis (phosphoenolpyruvate carboxykinase) and TCA cycle enzymes were upregulated in desiccated ST, these being expressed at FPKM values close to or higher than hydrated NST, and is the only peak in their expression in ST (Primary metabolism tab, Senescome; Fig. **S2**). This correlates with the observed increased accumulation of TCA cycle compounds malic, succinic and citric acid in desiccated ST relative to desiccated NST (Radermacher *et al*., 2019). There was also an increase in fermentation enzyme transcripts (L-lactate hydrogenase and aldehyde dehydrogenase) in desiccated leaves of all three tissues, but particularly in ST, indicating that fermentation may be employed to generate NADH. Another potential source of NADH might be glutamate dehydrogenase 2, upregulated in desiccated ST, which catalyzes the conversion of glutamate and NAD+ to 2-oxoglutarate and NADH (The UniProt consortium, 2019). Similarly, nitrate reductase, which catalyzes the conversion of nitrite and NAD+ to NADH and nitrate in the cytoplasm showed increased transcript abundance in desiccated ST. Alcohol dehydrogenase converts primary alcohols and NAD+ to an aldehyde and NADPH. The enzyme 2-alkenal reductase converts alkanals into alk-2-enals with high specificity for NADP+ and the release of NADPH (The UniProt consortium, 2019). This suggests that numerous sources for NAD(P)H generation exist despite the shutdown of photosynthesis and the Calvin cycle. The potential fate of this NAD(P)H is intriguing. The upregulation of ubiquinol oxidase 1a (AOX), an electron transfer flavoprotein subunit alpha and NADPH:quinone oxidoreductase in ST suggests that mitochondrial electron transport is somewhat functional as the cells approach the desiccated state. If translated, the AOX protein could compete for electron flow, suggesting that some electrons may be terminating at this electron acceptor (after passing through complexes I and II and generating some proton flow into the inner mitochondrial space) rather than proceeding through complex III towards complex IV via cytochrome C. The termination of electron flow at AOX rather than complex IV would dramatically reduce ATP yield (Vanlerberghe, 2013). There also seems to be some evidence that electron transport is still functioning in the near-desiccated state, as ATP synthase transcript abundance overall is not significantly different during desiccation to that at FT in NST (Primary metabolism tab, Desiccome) and there appears to be an increase in cytochrome c abundance in all tissues during all stages of water deficit (Primary metabolism tab, Senescome).

#### Transport: nitrogen and carbon

Integral to the process of senescence is the transport of breakdown products of metabolism from senescent to surviving tissues. Our study shows the significant accumulation of several nitrogen and carbon transporters in AD ST, suggesting that such transport is feasible.

Redistribution of nitrogen from source (senescent) to sink (non senescent) tissues is usually achieved by transport of amino acids, peptides and nitrate. Transcripts of several such transporters were significantly accumulated in the AD ST relative to other tissues (see Transport tab, Senescome). These included i) A tetra- and penta-peptide transporter, Oligopeptide transporter 7 (OPT7) which may play a role in transporting short peptides from senescing cells; ii) Cationic amino acid transporter 1 (CAT1), responsible for transport of a broad range of amino acids into and out of cells in Arabidopsis (Su et al., 2004); iii) Proline transporter 2, responsible for amino acid retrieval from the apoplast (Lehmann *et al*., 2011) and which in *X. schlechteri* could function as a potential transporter of proline from ST to NST/PST; iv) Protein NRT1/ PTR FAMILY 7.3, a transporter responsible for export of nitrate into xylem (Lin *et al*., 2008) and which could facilitate export of nitrate from ST; and v) protein NRT1/ PTR FAMILY 4.6, a nitrate bidirectional transporter which has been shown to also transport ABA (Corratgé-Faillie and Lacombe, 2017). Two additional mechanisms of nitrogen transport are possible. There was considerable accumulation of a cystinosin homologue in AD ST, which in mammalian cell lines and yeast is responsible for transporting cystine from lysosomes following protein hydrolysis (Kalatzis *et al*., 2001) but as yet has not been reported in plant tissues. There was also upregulation of a Urea-proton symporter DUR3, shown to regulate transport of urea from the apoplast to phloem in Arabidopsis (Bohner *et al*., 2015) (see Transport-amino acids/misc tab, Senescome App). Import of nitrogen to NST in turn could be facilitated by amino acid permease 1 (involved with amino acid transport into cells) which was highly accumulated in NST in the late stages of water deficit (see Transport tab, Desiccome App) but not differentially expressed in ST, and Protein NRT1/ PTR FAMILY 6.2, a nitrate transporter (Corratgé-Faillie and Lacombe, 2017) also highly expressed only in AD NST and PST (see Transport tab, Desiccome App).

As for nitrogen, a number of transcripts associated with carbon transport significantly accumulated in AD ST. These included sugar transport proteins 7 and 13, sugar/proton symporters involved in the uptake of apoplastic hexoses in response to biotic stress (Lemonnier *et al*., 2014) and two UDP-galactose transporters which might be implicated in transport of cell wall precursors and in turn drive RFO synthesis in surviving cells (Yobi *et al*., 2017) Two probable sugar phosphate translocators (homologs of At3g11320 and At5g25400) significantly accumulated in all three tissue types, indicating that sugars may also be transported in their phosphorylated form for delivery of both carbon and phosphate to sink tissues. A fatty acid export protein 4 (Li *et al*., 2015) was highly upregulated in ST during rehydration (see Transport, Desiccome), might be utilised to transport fatty acids to viable tissues during recovery (Transport-sugars/lipids/misc tab, Senescome App).

These data suggest nutrient transport is feasible, but given the timing of expression of transporters it is likely that most are translated only on rehydration, when most of the nutrient mobilization is likely to occur.

### Regulation of the senescence co-expression subnetwork

#### Expression of closely clustered senescence genes is directly and/or indirectly modulated by WRKY transcription factors

The senescence-specific region of the co-expression subnetwork was interrogated to identify putative regulators of senescence. Upstream regions of up to 1kb were obtained for this subset of genes (565 nodes) and subjected to DREME analysis to identify enriched transcription factor (TF) binding motifs in these sequences relative to the full network (3206 nodes). Two regions were significantly enriched, a GTCAAHS site and a CRCGTW site, in which the upstream regions predominantly contained both motifs (Fig. **6a**). TomTom identified potential motif-binding TFs using a plant transcription factor database (Jaspar plants 2018 non-redundant) (Supplementary data – tomtom_senescence_subnetwork files). The motifs were most frequently observed within 300bp of the transcriptional start sites of genes in the senescence-specific subnetwork (Supplementary **S3**).

**Figure 6:**
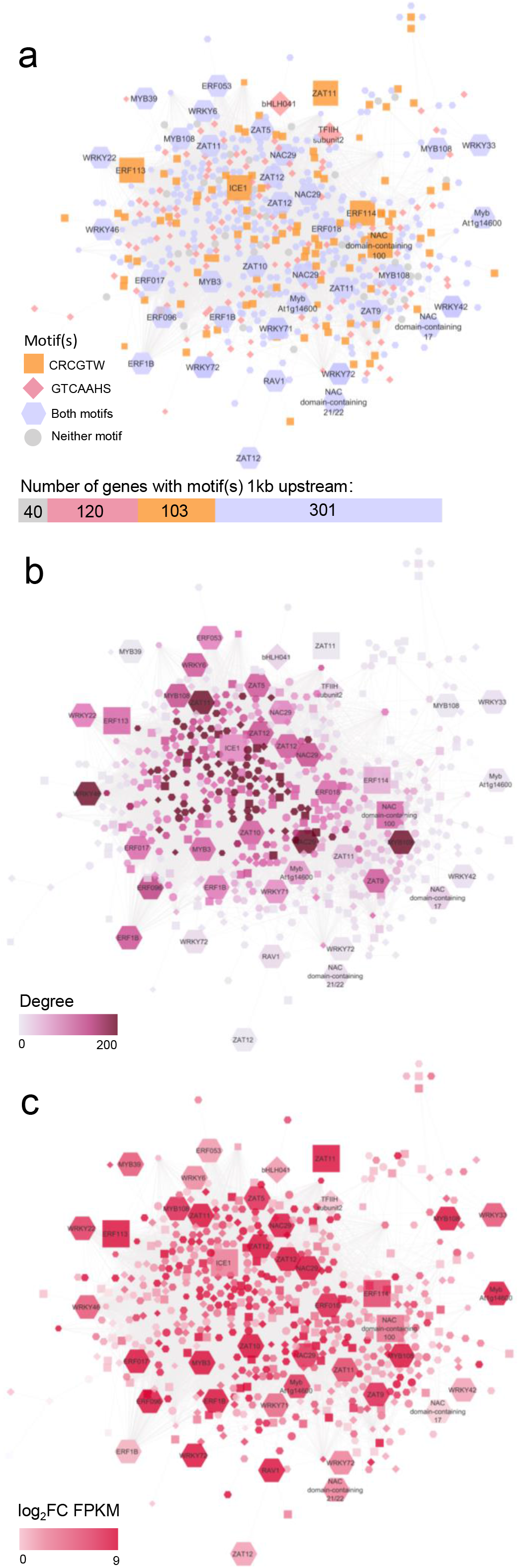
Transcriptional control of the senescence subnetwork. The senescence-specific subnetwork (564 nodes in total) was isolated from the main network and re-organized with the yFiles Organic layout to prevent node overlap. DREME analysis revealed enrichment of two motifs in the upstream region of genes in this subnetwork (GTCAAHS/ CRCGTW). The shape of nodes denotes whether their upstream regions contain the GTCAAHS motif (diamonds), the CRCGTW motif (squares), both the GTCAAHS and CRCGTW motifs (hexagons) or neither (circles). Transcription factor nodes are increased in size for greater clarity and labelled with their abbreviated SwissProt gene description. The nodes in panels A-C are coloured for their enriched motif (A), Degree (B) and log2FC in desiccated ST relative to hydrated NST (C).

Of the GTCAAHS motif-binding TFs, 15 were significantly upregulated in the Senescome, including four WRKY23, one WRKY57, one WRKY6, one WRKY70, two WRKY71 and three WRKY75 homologues (Supplementary data – tomtom_senescence_subnetwork_gtcaahs.xlsx), with eight also present in the network (Fig. **6**). Of the CRCGTW motif binding TF homologues, 9 were present in the ST-late response supercluster, but only one BHLH13 homologue (Supplementary data - tomtom_senescence_subnetwork_crcgtw.xlsx) was significantly upregulated in the Senescome. Figure **6a**, which denotes genes coloured based on the presence of these motifs in their upstream regions, shows their extensive presence and indeed how many genes in the senescence subnetwork are potentially under WRKY transcription factor control. All WRKY TFs present in the network contained one or both domains, indicating that their expression is self-regulated (Fig. **6a**, Fig. **S4**). Furthermore, most of the TFs in the senescence subnetwork contained these motifs in their upstream region, having a high degree of connectedness (Fig. **6b**) and expression in desiccated ST (Fig. **6c**), strongly suggesting a role for WRKY TFs in expression of this subnetwork.

#### A model predicting *Xs*SAG transcriptional regulation via MAPK6 activation of MYB3 and WRKY6

A potential mechanism for the control of the senescence-specific subnetwork is detailed in Figure 7. Having determined that many TFs in the subnetwork are potentially under control of the WRKY transcription factors, the upstream regions of WRKY TFs in the network were investigated more extensively using MEME. All eight WRKY TFs in the network possessed an ERF096 and/or ERF1B binding site in their upstream region. The ERF096 binding site was also found upstream of the ethylene biosynthetic genes ACC oxidase and ACC synthase and upstream of ERF096 itself (summarized in Fig. **S4**). ERF096 also had an upstream motif of a binding site for MYB3, which was also highly upregulated within the subnetwork. The ACC oxidase, MYB3 and all seven WRKY genes all contained a WRKY binding motif in their upstream regions, indicating a potential cyclical regulatory nature within this tightly associated subnetwork. Additionally, the ABA synthesis enzyme 9-cis-epoxycarotenoid dioxygenase NCED3 (chloroplastic) contained both motifs in its upstream region, indicating that ABA synthesis in late stages of drying may be driven by the action of this transcriptional regulatory network. Indeed, ABA is elevated in ST below 35% RWC, unlike in NST and PST, strongly suggesting a role for ABA-mediated responses in ST (Fig. **S5**).

**Figure 7:**
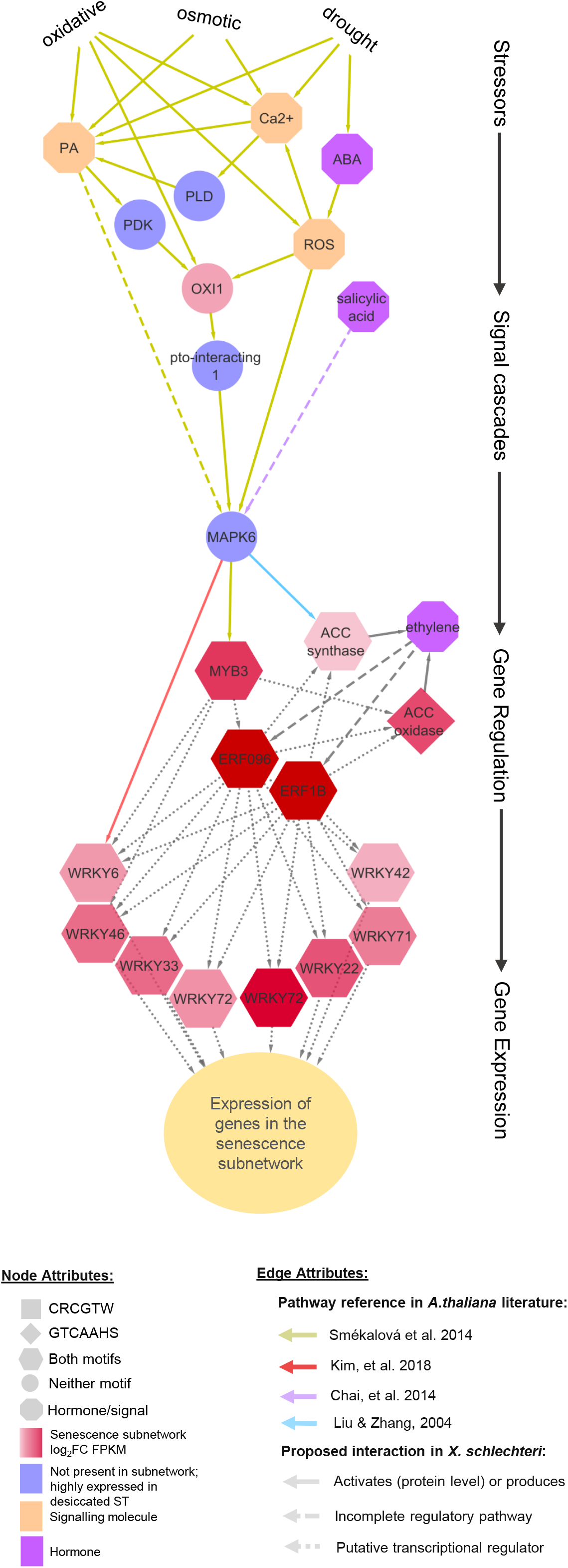
Proposed model for regulation of the senescence subnetwork based on studies in Arabidopsis. Nodes in shades of red are a subset of nodes from the senescence subnetwork proposed to play a role in signaling during senescence in X. schlechteri. Nodes are shaped based on the presence, combination absence of enriched motifs in their upstream regions. Severe abiotic stresses result in widely-reported elevations in stress signals (orange) phosphatidic acid (PA), intercellular calcium (Ca2+), abscisic acid (ABA), salicylic acid (SA) and reactive oxygen species (ROS). There are many targets of these signals (un-/understudied targets from senescence subnetwork shown in Supplementary figure 4), but of special interest to the current study is interaction with regulators up- and downstream from mitogen-activated protein kinase (MAPK)6, one of the studied regulators in senescence processes also upregulated in the Senescome. MAPK6, activated downstream of oxidative signal inducible 1 (OXI1) and through other mechanisms, has been shown to activate WRKY6 in senescent *A*.*thaliana* (reviewed in Kim et al. 2018). All 7 WRKY TFs in the senescence subnetwork contain ERF096 binding sites in their upstream regions (identified by TomTom analysis), indicating potential regulation via this TF in addition to MAPK6. ERF096 and ERF1B also potentially drive ACC oxidase and ACC synthase expression, as binding sites for both TFs are also present in their upstream regions. These gene products drive biosynthesis of ethylene, which in turn drives ERF096 and ERF1B expression, driving WRKY expression and thus regulation of the majority of genes within the senescence subnetwork. ACC synthase is additionally directly phosphorylated by MAPK6, leading to its activity in response to stress. Orange nodes represent nodes not present in the subnetwork but elevated in the Senescome. Acronyms: phospholipase D (PLD), 3-phosphoinositide-dependent protein kinase 1 (PDK).

We further propose that gene expression in the senescence subnetwork is reliant on activation of MYB3 and WRKYs to drive ethylene synthesis (Fig. **7**). This might occur via action of MAPK6 (highly expressed in Senescome: Signalling/Kinases tab), a known regulator of abiotic stress responses with a proposed role in senescence (Chai *et al*., 2014). The activity of signalling kinases is not necessarily related to the abundance of transcripts, as their regulation occurs at the protein level. Thus our speculation on their potential mechanisms is done cautiously. In Arabidopsis, drought, osmotic and oxidative stress result in the accumulation of calcium ions, ROS and phosphatidic acid within cells (Smékalová *et al*., 2014). Their accumulation is proposed to activate the serine/threonine receptor kinase OXI1 (highly transcriptionally upregulated in desiccated ST of *X. schlechteri* (Fig. **3**), which triggers MAPK6 activation (Rentel *et al*., 2004; Shumbe *et al*., 2016) and which in turn activates various transcription factors, including MYB3 (Smékalová *et al*., 2014) and WRKY6 (Kim *et al*., 2018). MAPK6 has been shown to directly phosphorylate ACC-synthase, driving stress induced production of ethylene in Arabidopsis (Liu and Zhang, 2004). The activation of WRKY6 has been shown to drive senescence-associated genes in Arabidopsis including calmodulins, peroxidases, pectin esterase, lipase, MAP kinases, serine/threonine kinases, ABC transporters and others (Robatzek and Somssich, 2002). We propose that a similar mechanism for ethylene production and WRKY TF synthesis is occurring in ST of *X. schlechteri*, via MAPK6 activation of MYB3 and WRKY6 (Fig. **7**). Collectively this data strongly indicates that processes comparable to stress-induced senescence in Arabidopsis is occurring in *X. schlechteri*, but at remarkably low water contents.

#### Regulation via the CRCGTW motif – a senescence repressor?

The enrichment of the CRCGTW motif in the upstream region of a substantial number of nodes, most of which also contain the GTCAAHS site, is also of interest (Fig. **6a**). TomTom analysis shows that this CRCGTW motif is a potential binding site for a number of TFs including bHLH13 and MYC2/3/4. Group IIId bHLH TFs (including bHLH13) have been shown to repress expression of senescence associated protease SAG29, whereas MYC3/4/5 TFs have been shown to bind to the same site and activate expression of this gene in the presence of jasmonic acid (Qi *et al*., 2015). Thus, repressed expression of these genes in NST and PST may be facilitated by bHLH TFs.

## Discussion

### Senescence occurs at extremely low water contents

This study shows that expression of XsSAGs occurs below the threshold of ca 40% RWC that is proposed to delineate vegetative desiccation tolerance and sensitivity (Zhang and Bartels, 2018; Pardo et al., 2020). Thus, senescence is occurring in context of inherent desiccation tolerance and, upon reaching 35% RWC, ST has transcribed all necessary components required to mount an effective defense against extreme water deficit. Furthermore, increase in transcription of senescence-associated genes below this water content, suggests ongoing select metabolic activity. Whether these are translated remains to be shown, but it is likely that many are. Both seeds and resurrection plants may accumulate long-lived mRNAs which are translated during rehydration (Sajeev et al., 2019; Farrant and Kruger, 2001). Molecular stabilization of transcriptional machinery is presumably required to facilitate the expression of XsSAGs and their regulators, and significant changes in the metabolite pool in these tissues (Radermacher et al., 2019) suggest some degree of ongoing metabolism at these extremely low water contents. The functionality of this mechanism can only be speculated upon, as at such water contents classical nutrient transfer to surviving tissues is unlikely. However it is possible that such transfer occurs on rehydration, with the few surviving cells bordering such tissues acting as conduits for transfer to the PST.

The observation that metabolic activity, in terms of transcription, translation, and metabolite changes can occur at RWCs below 30%, with respiration in many instances ceasing only at 10% RWC, is not new (reviewed in Costa *et al*., 2017b; Oliver *et al*., 2020). It has been postulated that this might be made possible due to formation of subcellular pockets in which molecular mobility is maintained in the absence of free or weakly bound water (du Toit et al., 2020; Farrant and Hilhorst, 2021). Such environments of mobility may include oil droplets, which remain fluid within dry cells, (Ballesteros *et al*., 2020), and liquid–liquid phase transitions, enabling molecular interactions, such as protein binding, in dry cells (Belott et al., 2020). Moreover, while difficult to demonstrate, it has recently been proposed that pockets of natural deep eutectic solvents (NaDES) may form during the late stages of dehydration in resurrection plants enabling ongoing select metabolism (du Toit et al., 2020; Farrant and Hilhorst, 2021). Indeed, the accumulation of NaDES forming metabolites citrate and sucrose at water contents below 35% RWC has been demonstrated in both NST and ST in *X. schlechteri* and it has been postulated that this could account of ongoing respiration observed at extremely low water contents (Radermacher et al., 2019). The slightly larger accumulation of malate and sucrose in ST could suggest an alternative source (or site) of NaDES formation. While highly speculative, it is possible that these do allow for ongoing and selectively different metabolisms in the respective tissues.

### Senescence initiation in response to prolonged and cumulative stress

The present study found that, at the transcriptional level, protective mechanisms are deployed in NST, PST and ST to the same degree: LEA, heat shock, sugar synthesis and anti-oxidant encoding genes are transcribed in all tissue types. Transcription below 35% RWC in the ST takes place in the context of a desiccated cell, as evidenced by the upregulation of desiccation tolerance genes. While clearly transcribed in ST, there is as yet no evidence as to whether they are translated and thus whether these gene products are performing their function in stabilising cells of the ST and ameliorating the stresses brought about by desiccation. Assuming they are translated and that their protein products are active, senescence is not the result of a deliberate switching off of desiccation tolerance processes (at the transcriptional level), rendering the tissue desiccation sensitive, but rather a parallel process engaging a relatively small number of regulators (WRKY, ERF and bHLH transcription factor families) to bring about macromolecular degradation, nutrient remobilisation and lysis in cells. The hypothesis that some of these Desiccome genes are indeed translated in senescent tissueis not unrealistic, as it is not unreasonable to assume that stabilisation of the cellular constituents required for transcription of senescence genes below 35% RWC is reliant on the presence of stabilising molecules (such as LEA proteins, molecular chaperones, and NaDES formation etc) to maintain the correct conformation of transcriptional machinery below 35% RWC.

The question of how the overlapping osmotic, drought and oxidative stresses that spell the demise of ST do not cause the same response in NST and PST during desiccation is an intriguing one. While there is some evidence that expression of the senescence subnetwork is triggered to a degree in these proximal tissues (expression of genes in cluster 3 and 6 do peak to a lesser degree in NST and PST than in ST, and there is increased log2 foldchange of the senescence subnetwork (Figs. **2**-**4**), indicating that some cells may indeed succumb to the stress.

Why so many cells within ST succumb, leading to bulk tissue death, may be due to prolonged exposure to drought stress. These tissues are the first tissues to experience water deficit (Radermacher *et al*., 2019) and have also endured prolonged exposure to UV radiation compared to their NST counterparts (due to their being the oldest part of the leaf). Therefore, the level of photoreactive damage they incur is presumably greater. They express similar levels of protectant associated transcripts, but this is possibly insufficient to mitigate the extent of damage incurred during drying, and senescence processes are initiated in response to the many overlapping stress signals.

### Redox status and aging promote desiccation-induced senescence in X. schlechteri

Assuming translation of genes transcribed in late stages of dehydration in ST, and that their products are active, senescence is not the result of a deliberate cessation of desiccation tolerance processes, rendering the tissue desiccation sensitive, but rather a precisely regulated process involving a relatively small number of regulators (WRKY and ERF) responsive to stress signals (ROS, ABA, Ca2+, etc.) (Fig. **7**). As further modelled in Figure 8, we propose that a tipping point is reached where cells of the ST, being older and thus exposed to environmental stresses for more prolonged periods, are no longer able to keep up with the energy demands of ROS mitigation, with ROS production converging with other signals upon MAPK6 (or its Xs homologue) to stimulate expression of XsSAGs via ethylene signalling. In this model, activity of the MAPK6 (cascade) is the determining factor for activation of senescence-inducing pathways, informed by the environment, e.g. through upstream receptor-like kinases (Kim et al., 2018), and aging-dependent ROS-homeostasis. We further postulate the existence of senescence repressors in surviving tissues, with MAPK6 (and other kinases) and the CRCGTW motif being potential targets for such repressors.

**Figure 8:**
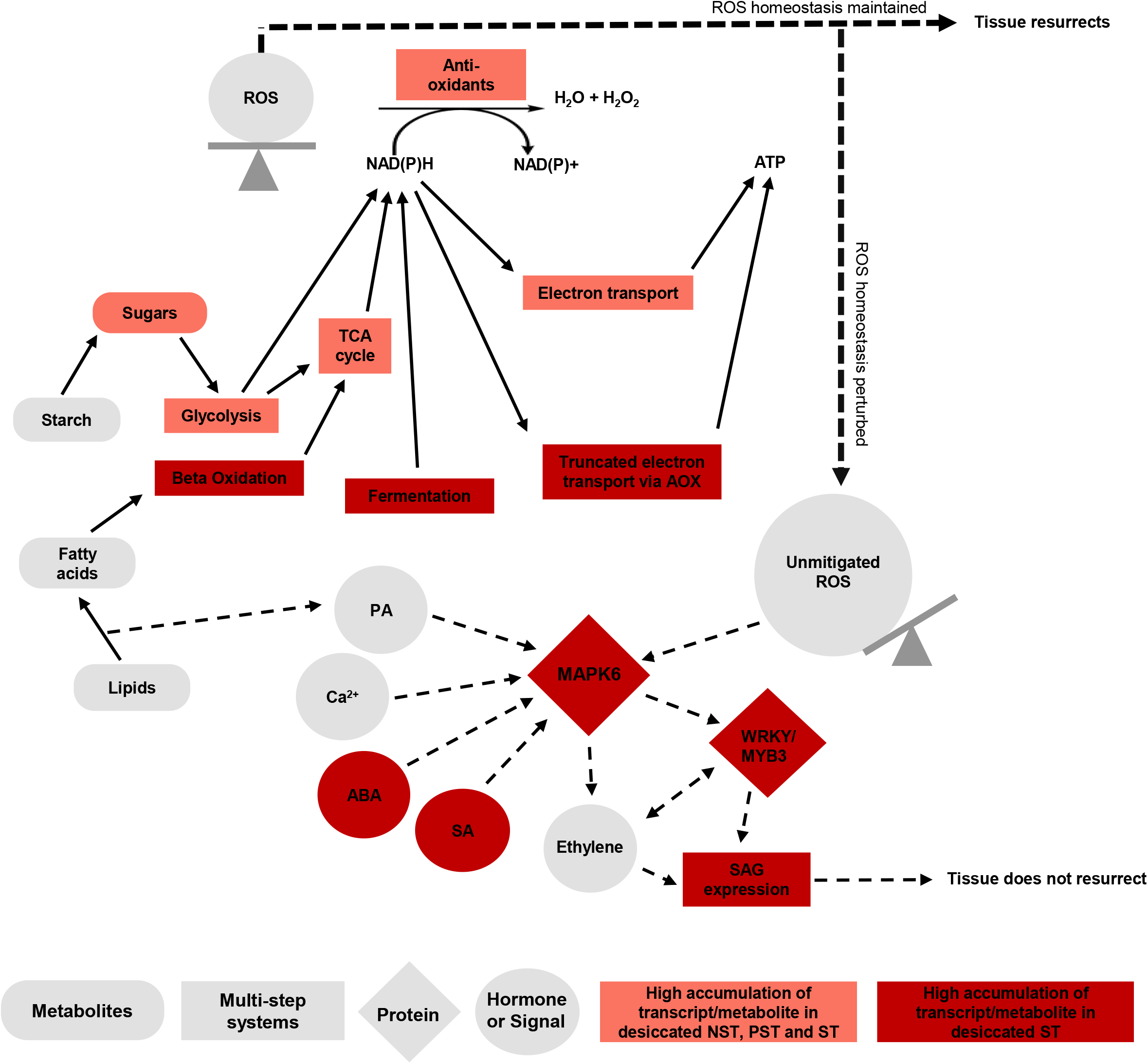
Hypothetical model coupling stress responses and energy requirements with senescence initiation in *X. schlechteri*. Stressed leaves shut down photosynthesis with the onset of water deficit. Glycolysis, the TCA cycle and electron transport in the mitochondria are presumably still active as transcripts encoding their enzymes do not diminish during desiccation, with some increasing in abundance (e.g. cytochrome C). These mechanisms drive production of ATP and NADH, required for a variety of intracellular protection mechanisms including production of vitrifying sugars and RFOs and mitigation of ROS via anti-oxidant enzymes (e.g. glutathione reductase) and for synthesis of non-enzymatic anti-oxidants (e.g. ascorbic acid). As the apical leaf tissues become severely stressed (<20% RWC) available sources for the regeneration of NADH and ATP become fewer. Expression of enzymes involved with beta-oxidation and fermentation increase in order to produce NADH, as does expression of alternative oxidase (AOX), which truncates the electron transport chain, terminating it short of cytochrome C. This truncation of electron transport potentially prevents production of free radicals while still producing some ATP for cellular processes. However, as energy sources for production of NADH and ATP dwindle, as does the cell’s ability to mount a defense against toxic intermediates and ROS. This accumulation of unmitigated ROS, coupled with phosphatidic acid build up (a result of the action of lipases required for entry of lipids into beta-oxidation), leads to triggering of senescence processes as described in figure 6 via MAPK6, and expression of senescence-associated genes (SAGs), leading to senescence as the cells approach the desiccated state.

Our model (Fig. **8**) proposes that redox balance plays an important role in this tipping point. The ability to retain antioxidant capacity is correlated with tissue longevity in the resurrection plant *Myrothanumnus flabellifolia* (Kranner et al., (2002) and it has been proposed that antioxidant potential, calculated as the gluthatione half-cell reduction potential ((EGSSG/2GSH) serves as a marker for cell viability and is a potential modulator of stress associated PCD in plants (Kranner et al., 2006). Although not measured in the current study, previous work has shown high redox potential in surviving tissues of this species (Farrant et al., 2007) and it is possible that this assists in suppression of senescence in such tissues.

### Senescence initiation in response to energy balance

A parallel argument can be made regarding the energy balance within cells. NADH and ATP are required by enzymatic anti-oxidants, such as glutathione reductase, and for synthesis of non-enzymatic anti-oxidants for dealing with ROS and free radicals generated during water deficit stress. Following the notion that the anti-oxidant status of a resurrection plant is the ultimate determinant of its viability we propose that to keep up with the demands for NADH and ATP, while limiting ROS formation, stressed cells shut down photosynthesis but keep glycolysis, the TCA cycle and electron transport active. As ST cells enter a severely stressed state, below 35% RWC, genes encoding lipases and beta-oxidation enzymes are transcribed to drive the TCA cycle for regeneration of NADH. The electron transport chain is truncated by AOX to limit high energy electron transitions and thus further unstable intermediate formation. Fermentation is additionally employed to meet energy demands. When a tipping point is reached, wherein the energy required to mitigate ROS and toxic intermediates is not produced rapidly enough to prevent their build-up, senescence processes are initiated via activation of ERF and WRKY genes as described in Figure 7. Additionally, the action of lipases to produce substrate for beta-oxidation, especially phospholipase C and D, drives production of phosphatidic acid, yet another driver for senescence initiation

The observation of increased expression of carbon and nitrogen transporters in AD ST and differential expression of uptake transporters in in NST and PST suggests a directional flow of nutrient from ST to NST/PST probably only on rehydration. Interestingly, like its sister species *Xerophyta humilis* (Meyers et al., 2010), *X. schlechteri* suppresses flowering during dehydration, with flowering occurring immediately after rehydration following a desiccation event (personal observation). We speculate that this could, in part, be related to remobilization of nutrients from senescent to surviving tissues during initial recovery and that it is furthermore possible that the traits of senescence- and flowering-repression are linked. Further investigation into this phenomenon raises the possibility of not only suppressing senescence in crops under drought, but altering flowering patterns during a drought so as to ensure flowering and thus crop yield despite of the drought.

## Supporting information

senescence_subnetwork_crcgtw

xerophyta schlechteri replicates fpkm

xerophyta schlechteri transcriptome data

fold change fischers exact test comparisons to ft nst degs

old change fischers exact test comparisons to ft nst

nonredundant GO terms

senescence subnetwork gtcaahs

DREME senescence subnetwork

## Acknowledgments

JMF acknowledges funding supplied by the South African Department of Science and Innovation and National Research Foundation (Grant No 98406).

## Author contributions

BW SGM and JMF conceived the study and mentored ALR who conducted the research did the analysis and wrote the paper. AI assisted in initial bioinformatic analysis and HD conducted the ABA analysis. HH provided critical insight to the interpretations of the work and JMF compiled the final manuscript.

## Data availability

Raw data is available through the corresponding author

## Supporting Information

**Fig. S1**. WRKY transcription factors are highly expressed in desiccated ST, while DNA remodellers are diminished.

**Fig. S2**. Transcription of central metabolism genes is most abundant below 35 RWC (sampling point 5 in apical ST).

**Fig. S3**. Frequency of TF binding sites.

**Fig. S4**. WRKY and ethylene responsive transcription factors may be co regulators of the Senescome.

**Fig. S5**. ABA is elevated in ST in the late stages of drying but not in NST/PST.

**Fig. S6**. qPCR validation of trends in the transcriptome

**Table S1**. An average of 87% of transcripts measured aligned to the reference genome.

## Supplementary

**Files:**

DREME_senescence_subnetwork.html

fold_change_fischers_exact_test_comparsions_to_ft_nst.csv

fold_change_fischers_exact_test_comparsions_to_ft_nst_degs.csv

nonredundent_GO_terms.xlsx

Supplementary data.Rproj

tomtom_senescence_subnetwork_crcgtw.xlsx

tomtom_senescence_subnetwork_gtcaahs.xlsx

xerophyta_schlechteri_replicates_fpkm.csv

xerophyta_schlechteri_transcriptome_data.csv

**Supplementary figure 1:**
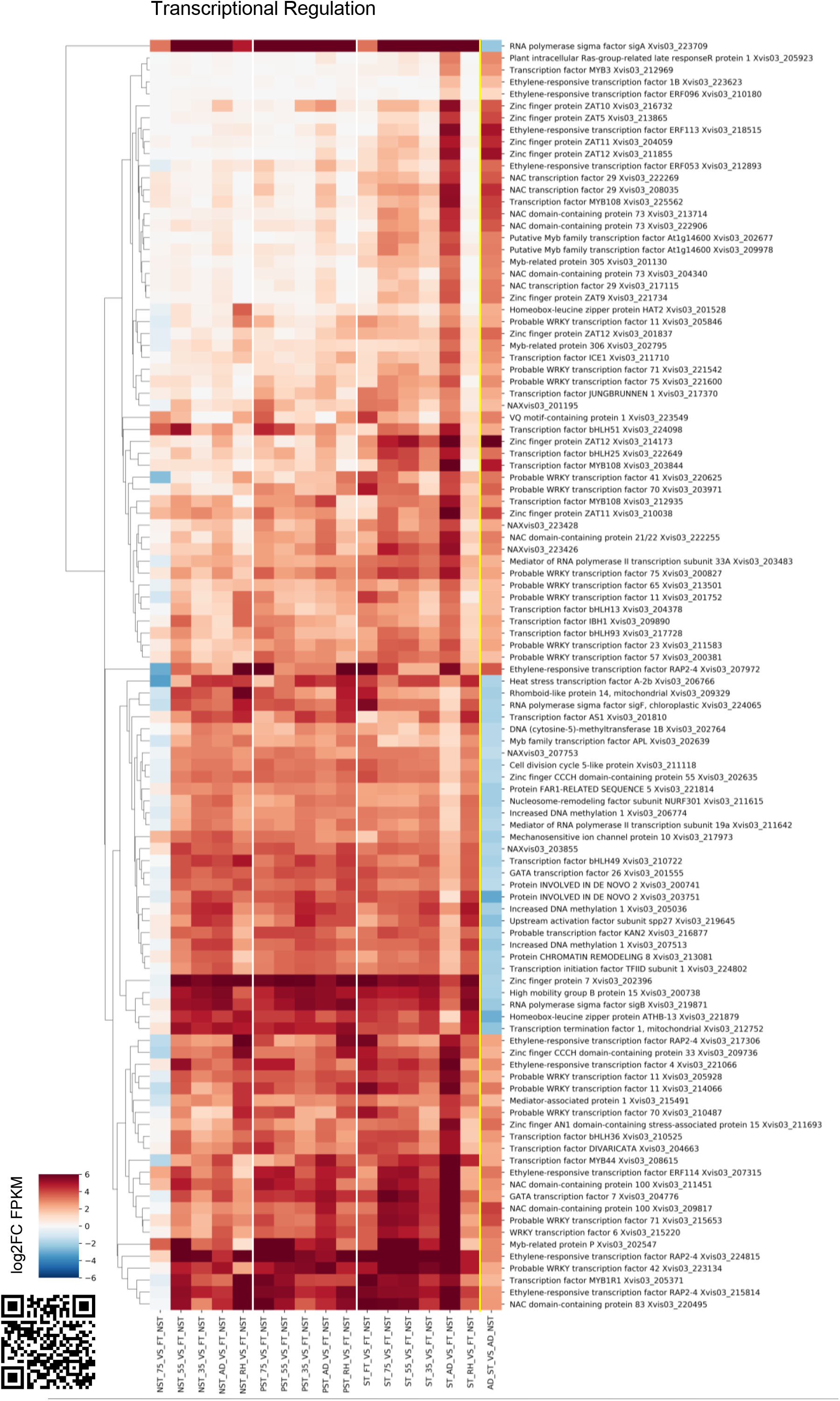
WRKY transcription factors are highly expressed in desiccated ST, while DNA remodellers are diminished. Senescence-specific DEGs encoding transcription factors were identified by comparing desiccated ST to desiccated NST. A subset of genes in the transcriptional regulation Mercator category are shown here, with their Log2 fold change values relative to FT NST shown in NST, PST and ST during water deficit and rehydration. Genes are labelled with their SwissProt annotation and genome ID. RWC values in the x-axis are representative of NST values. White vertical lines separate tissue types (NST-PST-ST). The right-most column represents gene expression in desiccated ST relative to desiccated NST. The full Senescome dataset is available at the QR code or https://astridite.shinyapps.io/XeroSenescome/

**Supplementary figure 2:**
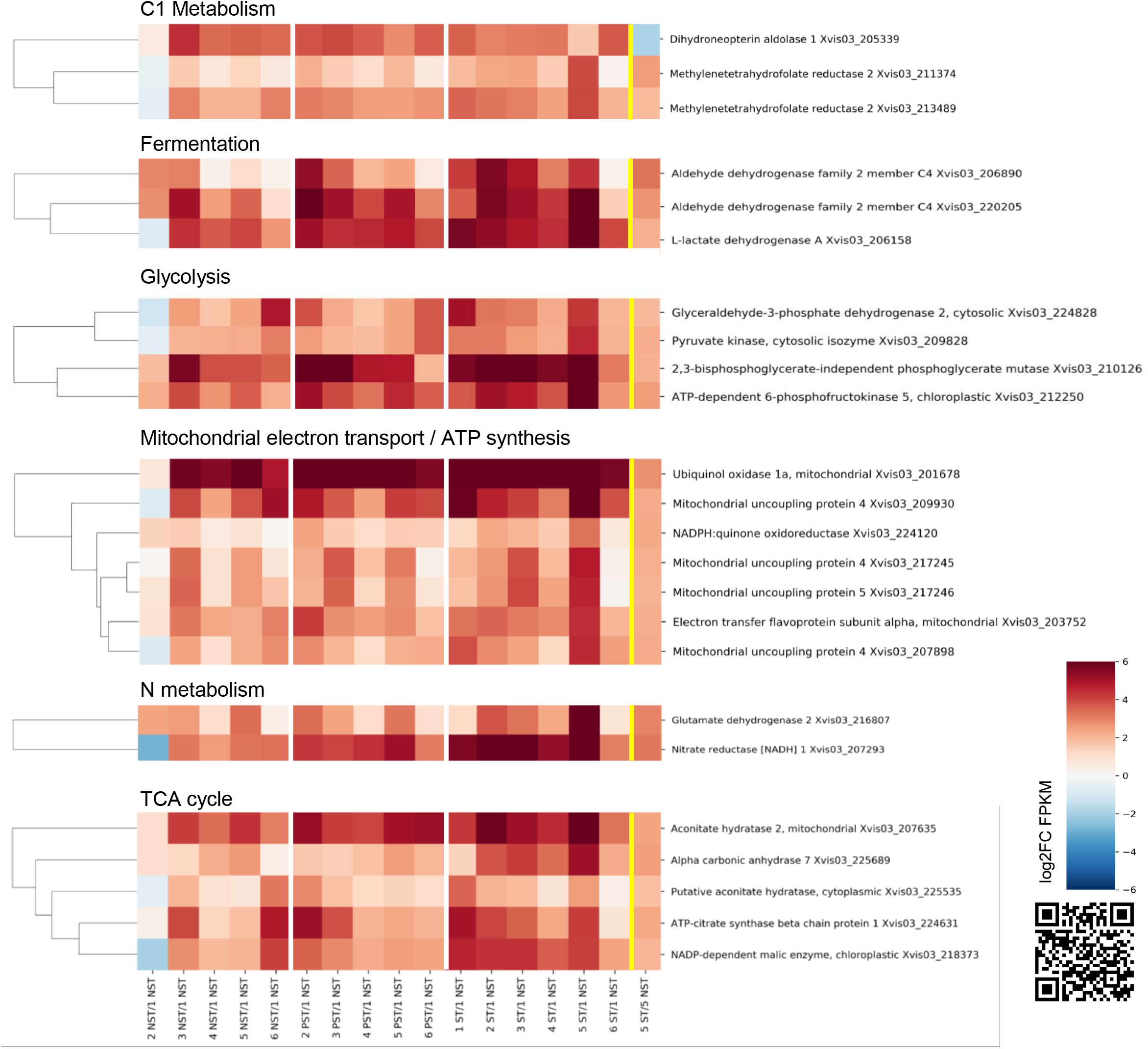
Transcription of central metabolism genes is most abundant below 35% RWC (sampling point 5) in apical ST. Senescence-specific DEGs were identified by comparing desiccated ST to desiccated NST. A subset of genes in the Mercator categories to do with central metabolism are shown here, with their Log2 fold change values relative to hydrated NST shown in NST, PST and ST during water deficit and rehydration. Genes are annotated with their SwissProt annotation and genome ID. X-axis labels are representative of the comparison, with the sampling point (number 1-6) and tissue type (NST/PST/ST) shown relative to hydrated NST (1 NST). White vertical lines separate tissue types (NST-PST-ST). The right-most column represents gene expression in desiccated ST relative to desiccated NST (5 ST/5 NST). The full Senescome dataset is available as a supplementary data set, or at the QR code (https://astridite.shinyapps.io/XeroSenescome/)

**Supplementary figure 3:**
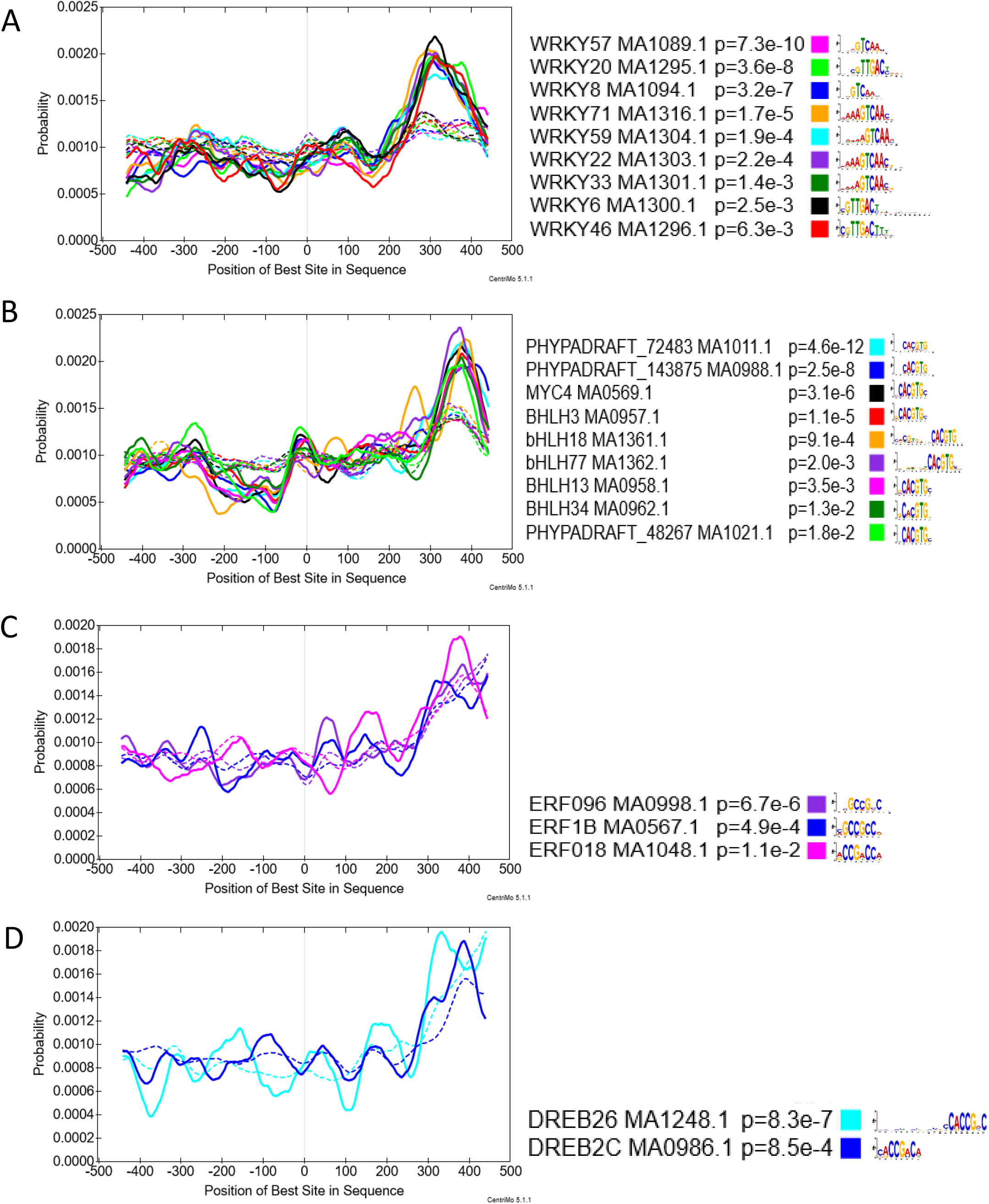
Frequency of TF binding sites. Centrimo analysis of upstream regions revealed that enriched transcription factor binding sites most frequently occur within 300bp from the transcriptional start site of the senescence subnetwork.

**Supplementary figure 4:**
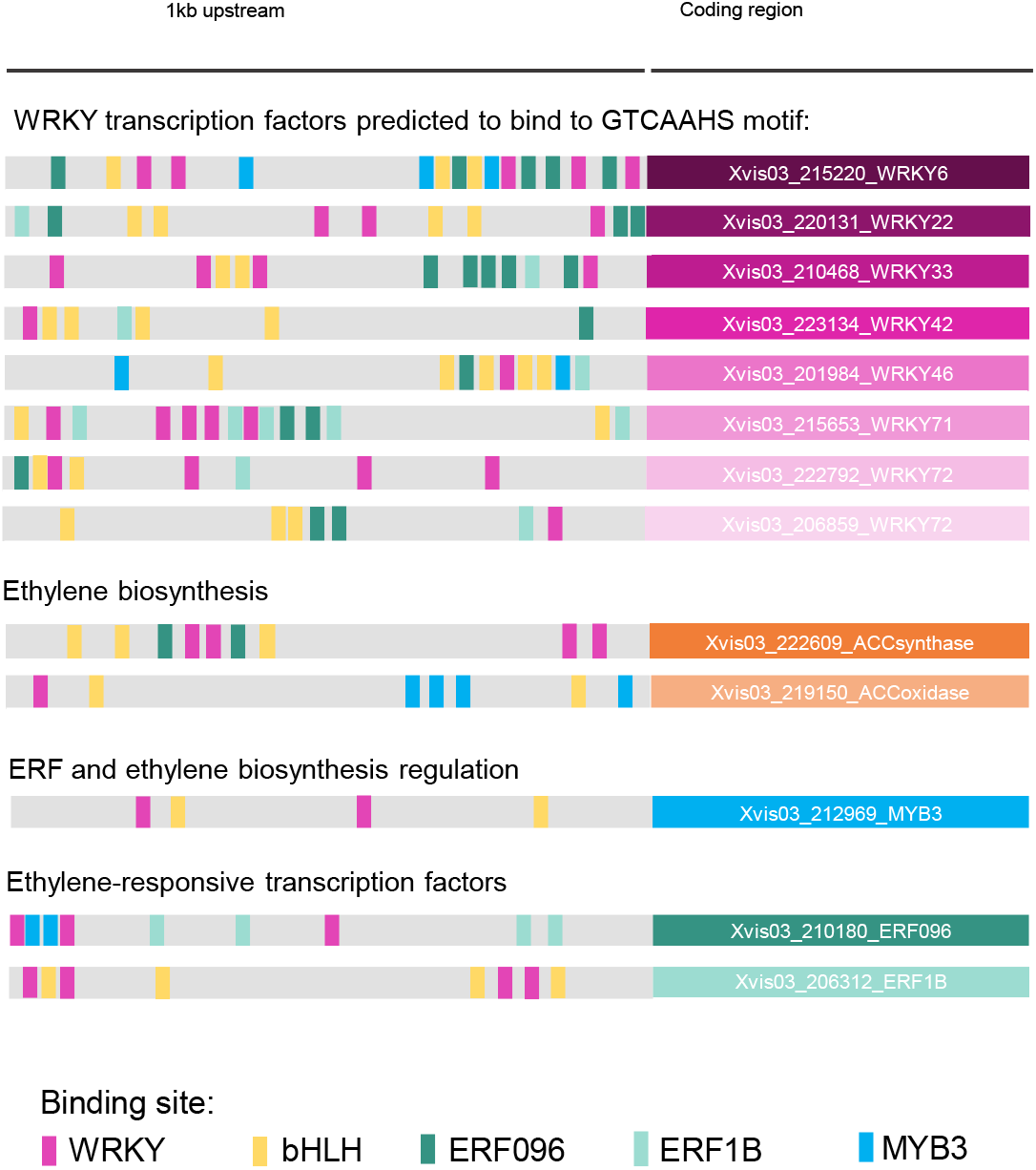
WRKY and ethylene-responsive transcription factors may be co-regulators of the Senescome. A summary of the TomTom/MEME upstream sequence analysis showing binding sites for specific regulators in predicted Senescome transcriptional regulators.

**Supplementary figure 5:**
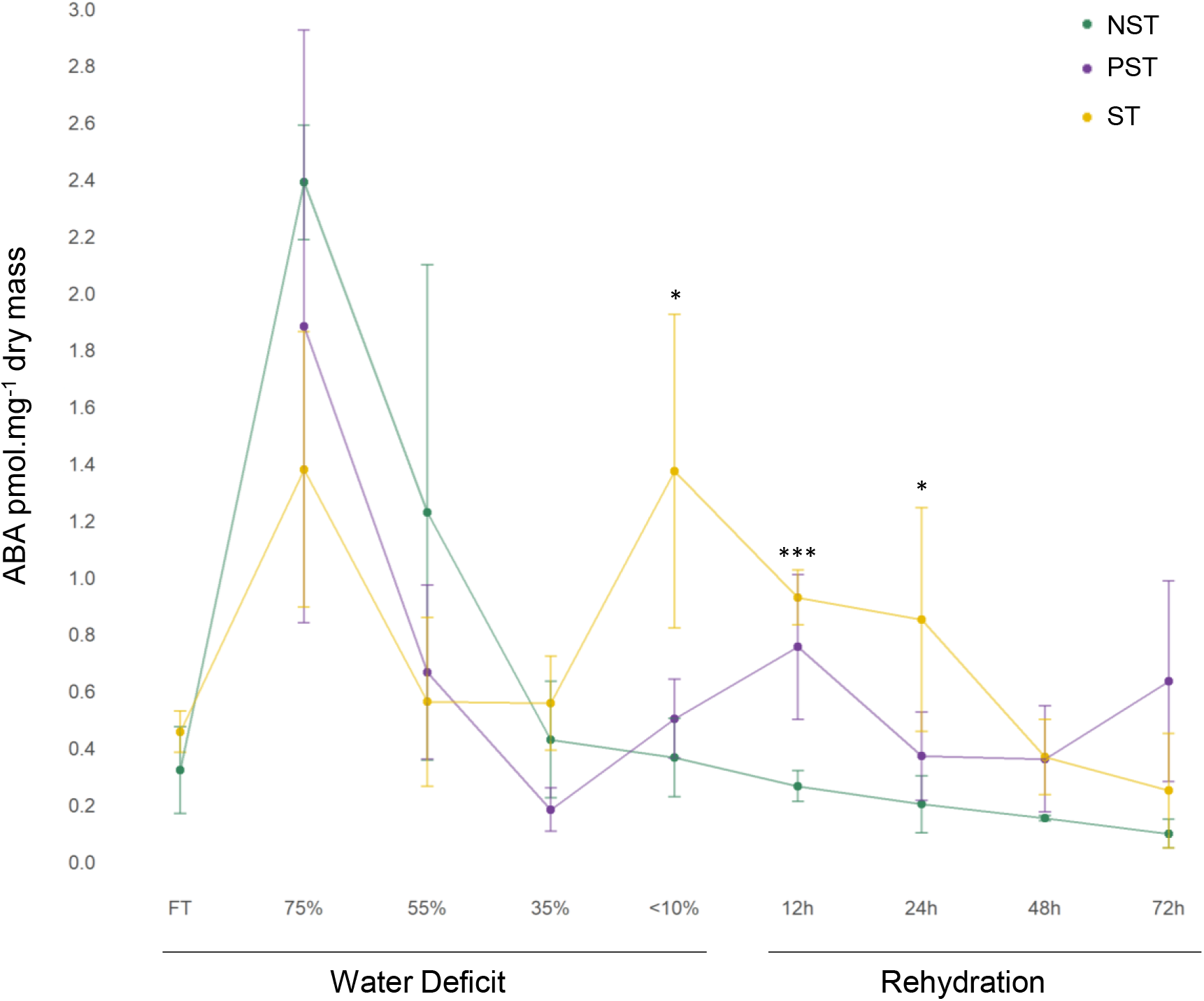
ABA is elevated in ST in the late stages of drying but not in NST/PST. ABA was quantified during dehydration and rehydration in NST, PST and ST. RWC values are those of NST. The mean and standard deviation of each sample group is given, with the results of a t-test for comparison between ST and NST. ABA content is only significantly increased in desiccated ST and in early rehydration (12h and 24h).

**Supplementary figure 6:**
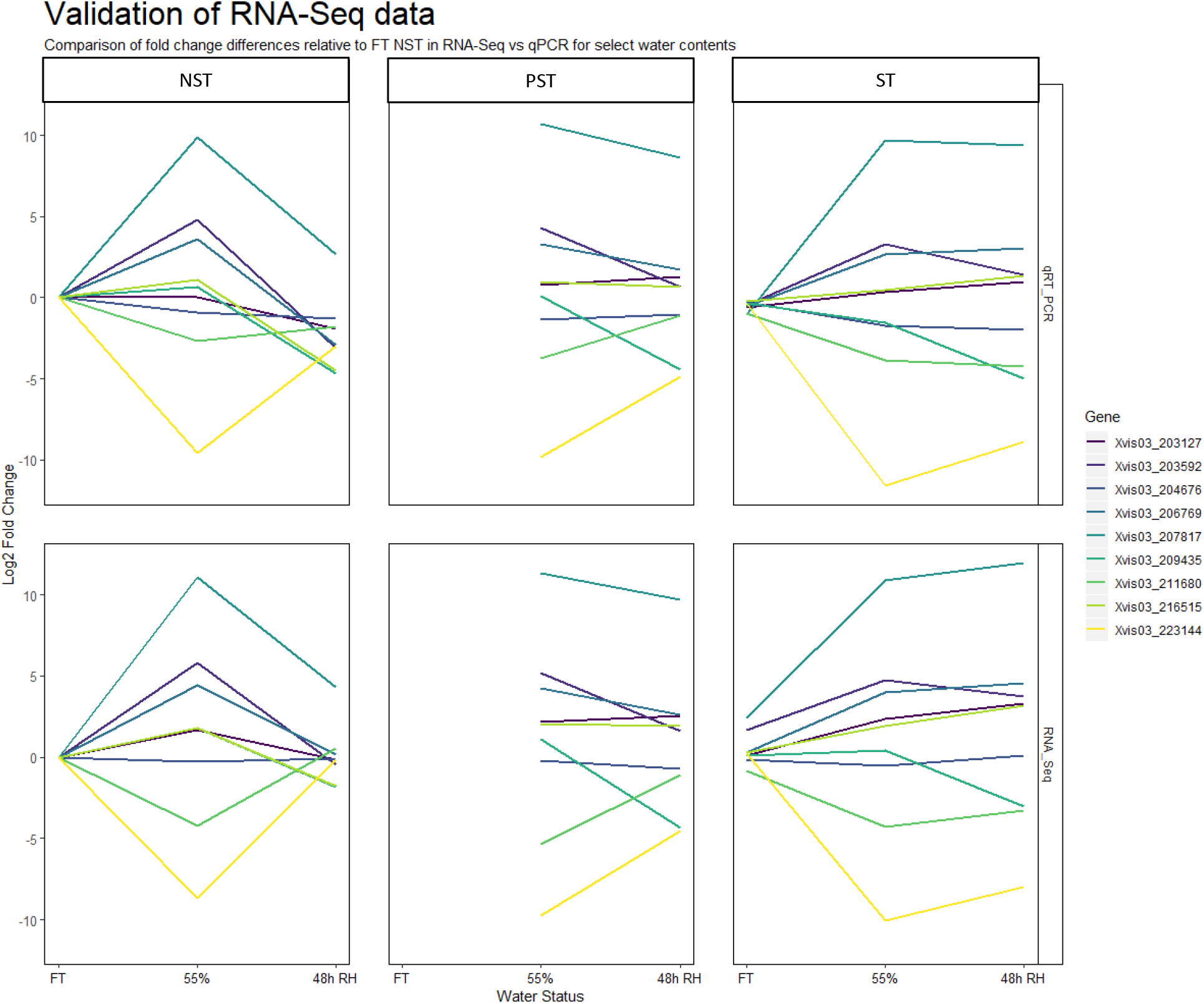
qPCR validation of trends in the transcriptome. RNA isolated from fully hydrated, 55% RWC (NST) and rehydrated samples was subjected to qPCR amplification to confirm consistency of the RNA Seq results.

**Supplementary Table 1:**
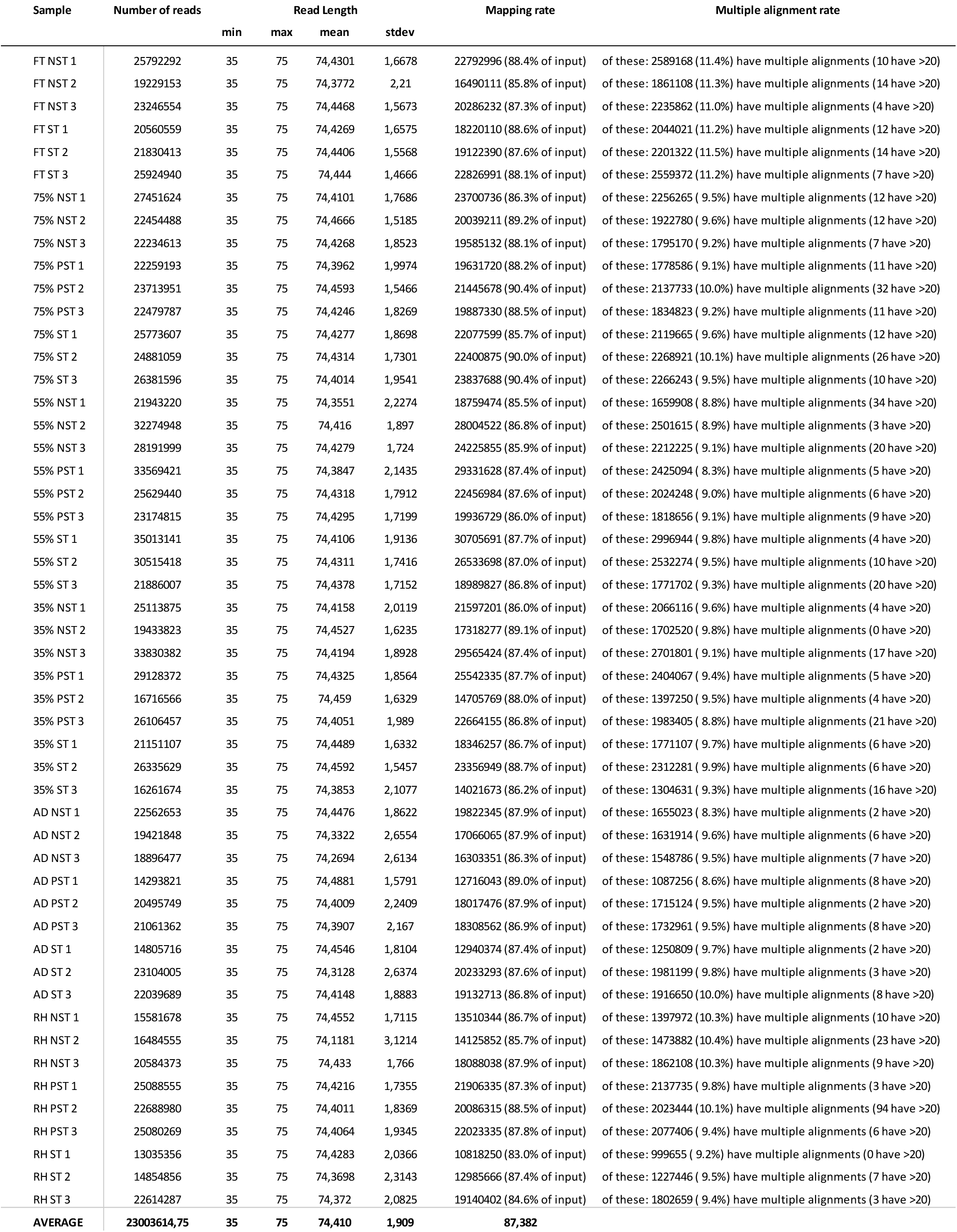
An average of 87% of transcripts measured aligned to the reference genome.

