## Supplementary material for "Desiccation-driven senescence and its repression in *Xerophyta schlechteri* are regulated at extremely low water contents": DREME senescence subnetwork: Suppl_Data_4_DREME_senescence_subnetwork.html


Help poup.

[close ]

DREME results in plain text format.

[
close ]

DREME results in XML format.

[
close ]

The name of the motif uses the IUPAC codes for nucleotides which has
a different letter to represent each of the 15 possible combinations.

The name is itself a representation of the motif though the position
weight matrix is not directly equivalent as it is generated from the
sites found that matched the letters given in the name.

Read more about the MEME suite's use of the IUPAC alphabets.

[close ]

The logo of the motif.

[close ]

The logo of the reverse complement motif.

[close ]

The E-value is the enrichment p-value times the number of candidate
motifs tested.

The enrichment p-value is calculated using Fisher's Exact Test for
enrichment of the motif in the positive sequences.

Note that the counts used in Fisher's Exact Test are made after
erasing sites that match previously found motifs.

[close ]

The E-value of the motif calculated without erasing the sites of
previously found motifs.

[close ]

Show more information on the motif.

[close ]

Submit your motif to another MEME Suite program or download your motif.

###### Supported Programs

Tomtom
:   Tomtom is a tool for searching for similar known motifs.

MAST
:   MAST is a tool for searching biological sequence databases for
    sequences that contain one or more of a group of known motifs.

FIMO
:   FIMO is a tool for searching biological sequence databases for
    sequences that contain one or more known motifs.

GOMo
:   GOMo is a tool for identifying possible roles (Gene Ontology
    terms) for DNA binding motifs.

SpaMo
:   SpaMo is a tool for inferring possible transcription factor
    complexes by finding motifs with enriched spacings.

[close ]

### positive sequences matching the motif / # positive sequences.

Note these counts are made after erasing sites that match previously
found motifs.

[close ]

### negative sequences matching the motif / # negative sequences.

Note these counts are made after erasing sites that match previously
found motifs.

[close ]

The p-value of Fisher's Exact Test for enrichment of the motif in
the positive sequences.

Note that the counts used in Fisher's Exact Test are made after
erasing sites that match previously found motifs.

[close ]

The E-value is the motif p-value times the number of candidate motifs
tested.

Note that the p-value was calculated with counts made after
erasing sites that match previously found motifs.

[close ]

The E-value of the motif calculated without erasing the sites of
previously found motifs.

[close ]

All words matching the motif whose uncorrected p-value is less than
0.01.

[close ]

### positive sequences with matches to the word / # positive sequences.

Note these counts are made after erasing sites that match previously
found motifs.

[close ]

### negative sequences with matches to the word / # negative sequences.

Note these counts are made after erasing sites that match previously
found motifs.

[close ]

The p-value of Fisher's Exact Test for enrichment of the word in
the positive sequences.

Note that the counts used in Fisher's Exact Test are made after
erasing sites that match previously found motifs.

[close ]

The word p-value times the number of candidates tested.

Note that the p-value was calculated with counts made after
erasing sites that match previously found motifs.

[close ]

The sequence file used by DREME to find the motifs.

[close ]

The alphabet of the sequences.

[close ]

The count of the sequences.

[close ]

The name of the alphabet symbol.

[close ]

The frequency of the alphabet symbol in the control dataset.

[close ]

###### Details

| Positives | Negatives | P-value | E-value | Unerased E-value |
| --- | --- | --- | --- | --- |
| / | / |  |  |  |

###### Enriched Matching Words

x

#### Submit or Download

⇧⬆

⇩⬇

Submit MotifDownload MotifDownload Logo

###### Submit to program

|  |  |  |
| --- | --- | --- |
|  | Tomtom | Find similar motifs in published libraries or a library you supply. |
|  | FIMO | Find motif occurrences in sequence data. |
|  | MAST | Rank sequences by affinity to groups of motifs. |
|  | GOMo | Identify possible roles (Gene Ontology terms) for motifs. |
|  | SpaMo | Find other motifs that are enriched at specific close spacings which might imply the existence of a complex. |

Format:

Count Matrix
Probability Matrix
Minimal MEME

|  |  |
| --- | --- |
| Format: | PNG (for web) EPS (for publication) |
| Orientation: | Normal Reverse Complement |
| Small Sample Correction: | Off On |
| Width: | cm |
| Height: | cm |

### DREME

#### Discriminative Regular Expression Motif Elicitation

For further information on how to interpret these results please access
http://meme-suite.org/.   
To get a copy of the MEME software please access
http://meme-suite.org.

If you use DREME in your research please cite the following paper:  

Timothy L. Bailey, "DREME: Motif discovery in transcription factor ChIP-seq data", *Bioinformatics*, **27**(12):1653-1659, 2011.
[full text]

Discovered Motifs
  |  
Inputs & Settings
  |  
Program Information
  |  
Results in Text Format 

 
  |  
Results in XML Format


### Javascript is required to view these results!

### Your browser does not support canvas!

#### Description

DREME senescence subnetwork vs full network

#### Discovered Motifs

Next Top

|  | Motif | Logo | RC Logo | E-value | Unerased E-value | More | Submit/Download |
| --- | --- | --- | --- | --- | --- | --- | --- |
| 1. | GTCAAHS |  |  | 1.2e-005 | 1.2e-005 | ↧↥ | ⇢ |
| Details Positives \| Negatives \| P-value \| E-value \| Unerased E-value; 291 / 565 \| 1200 / 3206 \| 2.8e-010 \| 1.2e-005 \| 1.2e-005 Enriched Matching Words Word  \| Positives  \| Negatives  \| P-value  \| E-value; GTCAAAG \| 87 / 565 \| 311 / 3206 \| 6.6e-005 \| 2.9e+000; GTCAAAC \| 91 / 565 \| 358 / 3206 \| 7.7e-004 \| 3.4e+001; GTCAACG \| 45 / 565 \| 156 / 3206 \| 2.6e-003 \| 1.1e+002; GTCAATG \| 62 / 565 \| 242 / 3206 \| 4.8e-003 \| 2.1e+002; GTCAACC \| 54 / 565 \| 206 / 3206 \| 5.7e-003 \| 2.5e+002; GTCAATC \| 56 / 565 \| 218 / 3206 \| 7.0e-003 \| 3.1e+002 | | | | | | | |
|  | Motif | Logo | RC Logo | E-value | Unerased E-value | More | Submit/Download |
| 2. | CRCGTW |  |  | 2.0e-003 | 8.2e-004 | ↧↥ | ⇢ |
| Details Positives \| Negatives \| P-value \| E-value \| Unerased E-value; 337 / 565 \| 1519 / 3206 \| 4.6e-008 \| 2.0e-003 \| 8.2e-004 Enriched Matching Words Word  \| Positives  \| Negatives  \| P-value  \| E-value; CACGTT \| 178 / 565 \| 686 / 3206 \| 2.0e-007 \| 8.8e-003; CGCGTA \| 70 / 565 \| 252 / 3206 \| 4.4e-004 \| 1.9e+001; CGCGTT \| 86 / 565 \| 365 / 3206 \| 6.9e-003 \| 3.0e+002; CACGTA \| 148 / 565 \| 689 / 3206 \| 8.3e-003 \| 3.6e+002 | | | | | | | |

#### Inputs & Settings

Previous Next Top

###### Sequences

| Source | Alphabet | Sequence Count |
| --- | --- | --- |
| senescenceomesubclusterupstreamregions.txt | DNA | 565 |

###### Control Sequences

| Source | Sequence Count |
| --- | --- |
| networkupstream.txt | 3206 |

###### Background

| Name | Bg. |  |  |  | Bg. | Name |
| --- | --- | --- | --- | --- | --- | --- |
| Adenine | 0.336 | A | ~ | T | 0.342 | Thymine |
| Cytosine | 0.167 | C | ~ | G | 0.155 | Guanine |

###### Other Settings

|  |  |
| --- | --- |
| Strand Handling | This alphabet only has one strand Only the given strand is processed Both the given and reverse complement strands are processed |
| # REs to Generalize | 100 |
| Shuffle Seed | 1 |
| E-value Threshold | 0.05 |
| Max Motif Count | No maximum motif count. |
| Max Run Time | 18000 seconds. |

Previous Top

###### DREME version

5.1.1
(Release date: Wed Jan 29 15:00:42 2020 -0800)

###### Command line
